# Splice-switching ASOs targeting an Alu-derived exon in the *AURKA* 5’UTR collapse an SRSF1-AURKA-MYC oncogenic circuit in pancreatic cancer

**DOI:** 10.1101/2025.04.14.648775

**Authors:** Alexander J. Kral, Lu Jia, GeunYoung Sim, Ledong Wan, Adrian R. Krainer

## Abstract

Pancreatic ductal adenocarcinoma (PDAC) remains a highly lethal malignancy, driven by oncogenic *KRAS* mutations and dysregulated oncogenes, such as *SRSF1*, *MYC*, and *AURKA*. Although KRAS-targeted therapies are in development, resistance mechanisms underscore the need to identify alternative vulnerabilities. Here, we uncover an SRSF1-AURKA-MYC oncogenic circuit, wherein SRSF1 regulates *AURKA* 5’UTR alternative splicing, enhancing AURKA protein expression; AURKA positively regulates SRSF1 and MYC post-translationally, independently of its kinase activity; and MYC in turn transcriptionally upregulates both *SRSF1* and *AURKA*. Elevated SRSF1 in tumor cells promotes inclusion of an exonized Alu exon in the *AURKA* 5’UTR, resulting in splicing-dependent mRNA accumulation and exon-junction-complex deposition. Modulating 5’UTR splicing with splice-switching antisense oligonucleotides (ASOs) collapses the oncogenic circuit, reducing PDAC cell viability and triggering apoptosis. Our findings identify *AURKA* alternative splicing as a critical regulatory node and highlight ASO-mediated splice-switching as a potential therapeutic strategy that simultaneously targets *SRSF1*, *AURKA*, and *MYC* oncogenes.

## INTRODUCTION

Pancreatic ductal adenocarcinoma (PDAC) is extremely lethal, with a 5-year survival rate of 10-13%.^1^ PDAC is driven by *KRAS* mutations, followed by inactivation of key tumor suppressor genes.^2–5^ In addition to mutational drivers, dysregulated expression of oncogenes, including *SRSF1,*^6^ *MYC,*^7,8^ and Aurora Kinase A (*AURKA*)^9–12^ further accelerates disease progression. However, targeting these oncogenic pathways remains challenging.

KRAS-targeted therapies, such as KRAS^G12C^ inhibitors, are approved for non-small cell lung cancer (NSCLC) but are ineffective for most PDAC patients, due to the low prevalence of KRAS^G12C^ mutations.^13^ KRAS^G12D^ inhibitors, currently in clinical trials, may offer a broader treatment option, but early studies have raised concerns about resistance mechanisms, like those observed with KRAS^G12C^ inhibitors.^13–19^ It is critical to identify additional pathways that PDAC cells use to evade KRAS inhibition.

We previously reported that the splicing factor SRSF1 promotes PDAC progression.^6^ We found that SRSF1 enhances MAPK signaling through alternative splicing (AS) of *IL1R1*, promoting tumor growth. *SRSF1* expression is upregulated via MYC-driven transcriptional activation. MYC, a potent oncogenic transcription factor,^20^ is frequently amplified in PDAC and is linked to KRAS-inhibitor resistance.^21^ Both MYC and SRSF1 remain difficult therapeutic targets, due to their pleiotropic roles and lack of conventional druggable domains. Developing a strategy to simultaneously disrupt these oncogenic networks could provide meaningful therapeutic benefits.

AURKA is an essential miotic serine/threonine kinase required for cell-cycle progression and frequently overexpressed in cancers, including PDAC.^12,22,23^ AURKA has two paralogs: Aurora kinase B (AURKB) and Aurora kinase C (AURKC). AURKB is also frequently overexpressed in cancer and promotes tumorigenesis, whereas AURKC is primarily involved in spermatogenesis.^24^ AURKA overexpression is sufficient to drive cellular transformation^25^, and AURKA has been a highly sought-after drug target, leading to the development of numerous kinase inhibitors.^26^ Despite decades of effort, such inhibitors have shown limited clinical efficacy, likely reflecting AURKA’s nuclear kinase-independent functions, which extend beyond its role in mitosis.^27^ Notably, AURKA is a transcriptional target of MYC,^28,29^ and stabilizes MYC and N-MYC by preventing their proteasomal degradation in various cancer contexts.^30,31^ These findings suggest that alternative approaches beyond kinase inhibition, such as direct *AURKA* knockdown, could be more therapeutically effective.^27,32,33^ Supporting this notion, *AURKA* knockdown in HeLa cells reduces SRSF1 protein expression post-translationally,^34^ implying that targeting AURKA at the protein level could simultaneously disrupt multiple oncogenic networks by impairing the expression and stability of both SRSF1 and MYC.

*AURKA* alternative splicing (AS) has emerged as an important regulatory mechanism, particularly splicing within its 5’ untranslated region (5’UTR), which is suspected to influence AURKA protein expression.^32,35^ *AURKA* exon 2 inclusion is more prevalent in tumors and potentially enhances translation, contributing to increased AURKA protein levels in cancer.^36,37^ Our previous work demonstrated that SRSF1 regulates numerous AS events in PDAC organoids^6^; re-analyzing these data, we identified an exon-inclusion event in the mouse *Aurka* 5’UTR, suggesting that SRSF1 can regulate AURKA protein levels through AS.

Based on these published results and our own findings, we hypothesized that SRSF1, AURKA, and MYC form a potent, multilayered, oncogenic, positive-feedback loop, wherein MYC transcriptionally activates *SRSF1* and *AURKA*, SRSF1 drives increased *AURKA* expression through AS, and AURKA stabilizes MYC in the nucleus and regulates SRSF1 post-translationally, thereby perpetuating the cycle.

In this study, we describe how SRSF1 regulates *AURKA* 5’UTR AS, how the resulting isoforms enhance AURKA protein expression through a splicing-dependent mechanism, and how the SRSF1-AURKA-MYC circuit operates. Translating these mechanistic findings, we developed antisense oligonucleotides (ASOs) that target *AURKA* AS as a strategy to therapeutically disrupt this oncogenic circuit in cancer.

## RESULTS

### SRSF1 regulates mouse *Aurka* 5’UTR splicing

In our previous work,^6^ we utilized mouse PDAC organoids from KSC *(LSL-**K**ras^G12D/+^; tetO-**S**rsf1; LSL-rtTA; Pdx1-**C**re*) mice to identify SRSF1-regulated AS events (Figure 1A). One of these events involves *Aurka* and is characterized in the present study (Figure 1B). Doxycycline-induced SRSF1 overexpression resulted in inclusion of a previously unannotated 791-nt exon in the mouse *Aurka* (*mAurka*) 5’UTR (Figure 1C). This SRSF1-dependent splicing change was highly significant (ΔPSI +37, p = 8.44E-07) and occurred independently of *Kras* mutational status.

**Figure 1.**
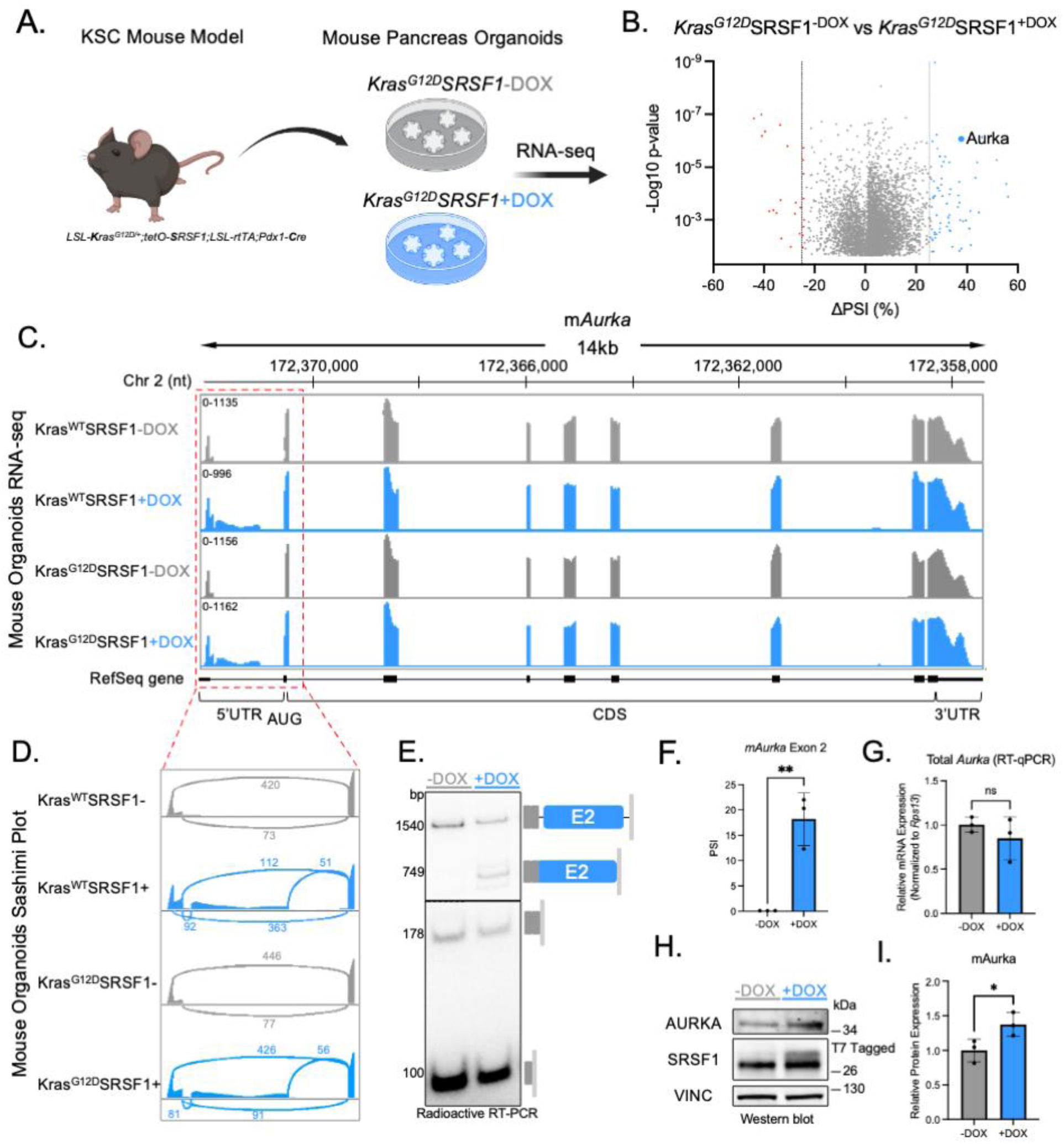
SRSF1 regulates mouse *Aurka* 5’UTR splicing (A) Workflow for generating KSC organoids for RNA-seq.^6^ (B) Volcano plot showing SRSF1-regulated splicing changes in KSC organoids. Red and blue dots represent AS events ≥ |25%| PSI and p ≤ 0.05. (C) IGV snapshot of the mouse *Aurka* gene. The range of number of reads is in the top left-hand corner. (D) Sashimi plot of the *Aurka* 5’UTR. Splice-junction reads are indicated. (E) Radioactive RT-PCR of *Aurka* 5’UTR from KSC organoids ±DOX (2 μg/ml for 48 h). (F) PSI quantification of exon 2 from (E) (n = 3 biological replicates). PSI, percent spliced in. (G) RT-qPCR of total *Aurka* from (E) normalized to *Rps13* (n = 3 biological replicates). (H) Western blot of AURKA, SRSF1, and Vinculin (VINC) from KSC organoids ±DOX. (I) Relative protein expression of AURKA compared to -DOX. Normalized to VINC loading control (n = 3 biological replicates). Data shown as mean±SD. ns = non-significant, *p ≤ 0.05, **p ≤ 0.01 by Student’s t test (F,G, and I).

A Sashimi plot^38^ of the *mAurka* 5’UTR revealed that it comprises two exons (exon 1 and exon 2) plus 5 nt of exon 3, preceding the AUG start codon. Exon 1 has two alternative 5’ splice sites (A5’ss), which correspond to exon 1a (98 nt) and exon 1b (176 nt). Under baseline conditions, the major isoform is exon 1a–exon 3. Upon SRSF1 overexpression, the exon 1b–exon 2–exon 3 isoform becomes expressed, incorporating the newly identified exon 2 (Figure 1D).

To confirm this AS event, we induced slight SRSF1 overexpression in the dox-inducible KSC organoids, and performed radioactive RT-PCR. Exon 2 inclusion was evident only upon SRSF1 induction, consistent with the RNA-seq analysis (Figure 1E). Total *mAurka* mRNA levels remained unchanged (Figure 1G), but AURKA protein levels increased, suggesting that exon 2 inclusion somehow enhances protein expression (Figure 1H-I). These findings demonstrate that SRSF1 regulates mAURKA protein levels by modulating 5’UTR AS.

### The human *AURKA* 5’UTR comprises two exonized Alu-elements that are upregulated in patient**-** derived PDAC organoids

Next, we investigated whether this AS event in the 5’UTR is conserved between *mAurka* and human *AURKA* (*hAURKA*). These orthologs share a conserved coding sequence (CDS) structure, comprising eight constitutively spliced exons. However, their 5’UTRs undergo AS, generating diverse mRNA isoforms that lack sequence conservation (Figure S1A-B). Alignment of mouse, chimp, rhesus, pig, and dog 5’UTR sequences to the *hAURKA* 5’UTR revealed sparse conservation outside the first 5’UTR exon. However, SRSF1 exonic splicing enhancer (ESE) motifs are partially conserved and scattered along mouse and human exon 2 (Figure S1C).

We found that the limited sequence conservation between human and mouse *AURKA* 5’UTRs reflects the presence of three overlapping Alu elements in human exons 2 and 3 (Figure 2A). AluSq2 (305 nt) fully overlaps exon 3, whereas AluSp (301 nt) and AluJo (66 nt) span exon 2 and its 5’ and 3’ splice sites (5’ss, 3’ss), respectively. Alu elements are primate-specific transposable elements that can undergo exonization by accumulating mutations that generate splice sites, increasing transcript diversity.^39–42^ Additionally, oppositely oriented Alu elements within 2 kb can form long double-stranded RNA (dsRNA) secondary structures, influencing splicing and other RNA-processing steps.^43–48^

**Figure 2.**
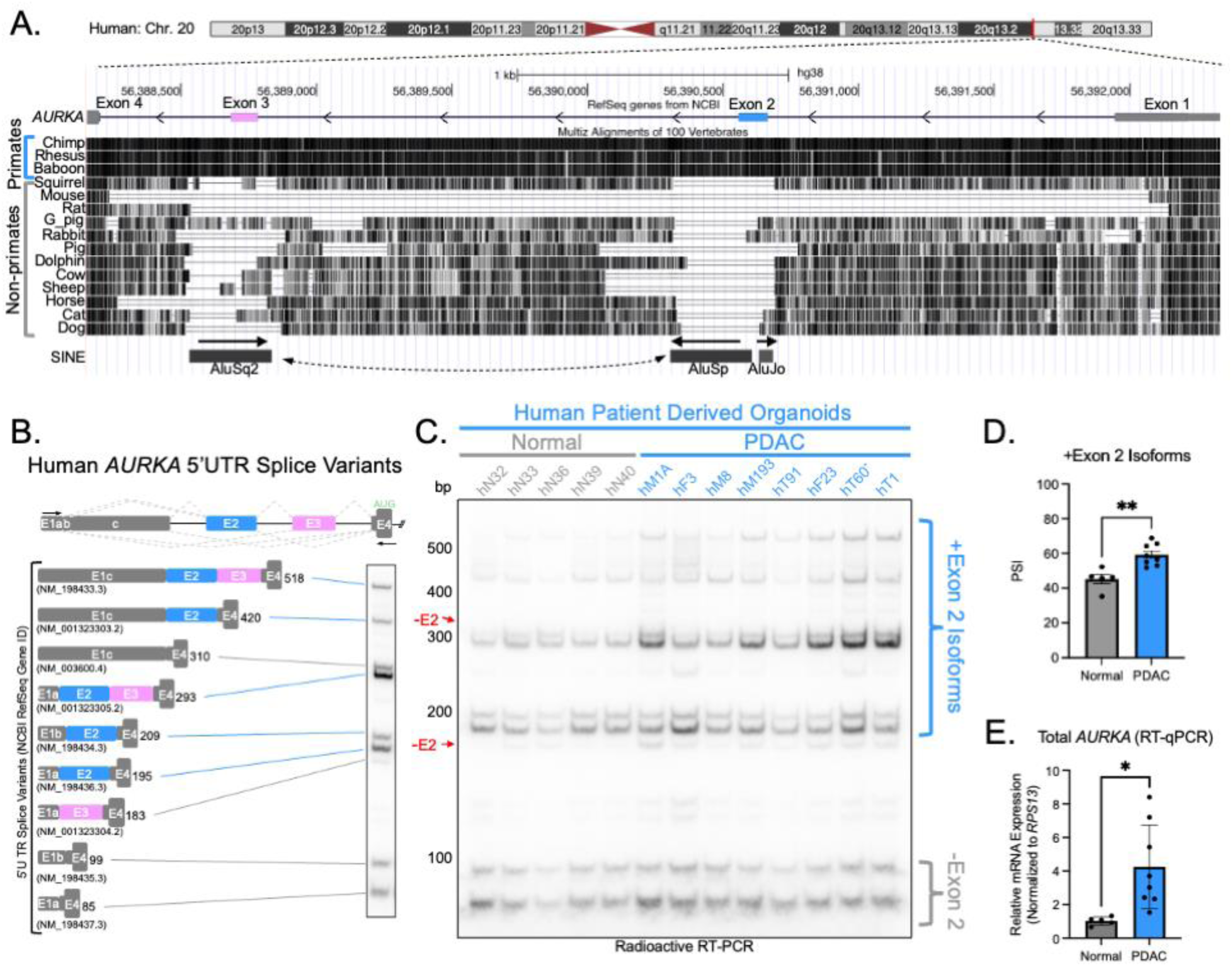
The human *AURKA* 5’UTR comprises two exonized Alu-elements that are upregulated in patient-derived PDAC organoids (A) UCSC genome browser view (https://genome.ucsc.edu/s/krala/AURKA_5UTR) of the conservation of the *AURKA* 5’UTR locus. Black lines = conserved nucleotide compared to human. AluSq2, AluSp, and AluJo elements are indicated at the bottom. SINE, short interspersed nuclear element. (B) Schematic of human *AURKA* 5’UTR gene architecture and splice variants. RT-PCR primers are indicated. Diagram of all observable RT-PCR product (with NCBI RefSeq mRNA accession numbers) and representative radioactive RT-PCR gel image (right). (C) Radioactive RT-PCR of *AURKA* 5’UTR from normal and PDAC patient-derived organoids. (D) PSI quantification of +E2 isoforms from (C). (E) RT-qPCR of total *AURKA* from normal and PDAC PDO lines normalized to *RPS13*. Data shown as mean±SD. *p ≤ 0.05, **p ≤ 0.01 by Student’s t test (D and E). See also Figures S1-S4.

AluSq2 and AluSp are nearly perfect reverse complements (Figure S2). RNAfold^49^ predicted that these elements can form a ∼300-nt dsRNA structure, potentially affecting splicing (Figure S3A-B). To infer the exonization event’s evolutionary history, we compared the splice sites of exons 2 and 3 to the ancestral AluS and AluJ sequences.^40,41,50,51^ Exon 3 fully overlaps AluSq2, with strengthened splice sites (MAXENT 3.61 → 11.44 for the 3’ss and 1.35 → 6.60 for the 5’ss), consistent with known Alu exonization patterns (Figure S3C).^39^ Exon 2 represents a chimeric exonization event, with AluSp providing the 5’ss (MAXENT - 1.44 → 7.07) and AluJo the 3’ss (unchanged; MAXENT 2.41) (Figure S3C). To our knowledge, this is the first report identifying *AURKA* exons 2 and 3 as exonized Alu elements.

The human *AURKA* 5’UTR undergoes extensive AS, with nine major and four minor isoforms annotated in the NCBI database. Among these are the exon-2-included (+E2) isoforms, which were reported to be preferentially expressed in cancer; however, their role in regulating AURKA protein expression remains unclear, and no reports have linked the +E2 isoforms to PDAC.^32,35–37^ Our analysis of the GTEx database demonstrates that the -E2 isoform, NM_198437.3, is the predominant isoform across all normal tissues (Figure S1D). +E2 isoforms are expressed in the testes and immortalized cell lines, though NM_198437.3 remains the major transcript.

To investigate the *AURKA* 5’UTR isoforms and the role of exon 2 in PDAC, we analyzed these mRNA isoforms using radioactive RT-PCR. We detected all nine major isoforms, with five +E2 variants identified and assigned to their respective bands (Figure 2B). Exon 1 has three A5’ss, corresponding to exon 1a (48- nt), exon 1b (+14-nt), and exon 1c (+ 211-nt). Many of the isoforms differ only in exon 1a/1b A5’ss usage, giving rise to doublets on gels. To assess PDAC-dependent AS changes, we examined *AURKA* 5’UTR isoforms across five normal and eight PDAC patient-derived organoid lines, revealing preferential exon 2 inclusion in PDAC, with an average +15% change in percent-spliced-in (ΔPSI) (Figure 2C-D). Total *AURKA* mRNA levels were also elevated almost 4-fold, on average, in the PDAC organoids, consistent with public RNA-seq data showing *AURKA* overexpression in PDAC tumors and a correlation with poor survival outcomes (Figure 2E and Figure S4A-B).

Taken together, these findings indicate that: i) *AURKA* exons 2 and 3 evolved as exonized Alu elements; and ii) the +E2 isoforms are upregulated in PDAC.

### SRSF1 regulates the *hAURKA* +E2 isoforms in PDAC organoids

We next investigated whether SRSF1 directly regulates *AURKA* 5’UTR splicing in human PDAC. Using the hT1 organoid line, we introduced a doxycycline-inducible SRSF1 cDNA construct via lentiviral transduction. After 48 h of doxycycline treatment, we assessed *AURKA* 5’UTR splicing and AURKA protein levels. Slight SRSF1 overexpression was sufficient to promote exon 2 inclusion, showing that SRSF1 regulates these isoforms (Figure 3A-B). Additionally, SRSF1 overexpression increased total *AURKA* mRNA and protein levels almost 2-fold (Figure 3C-E).

**Figure 3.**
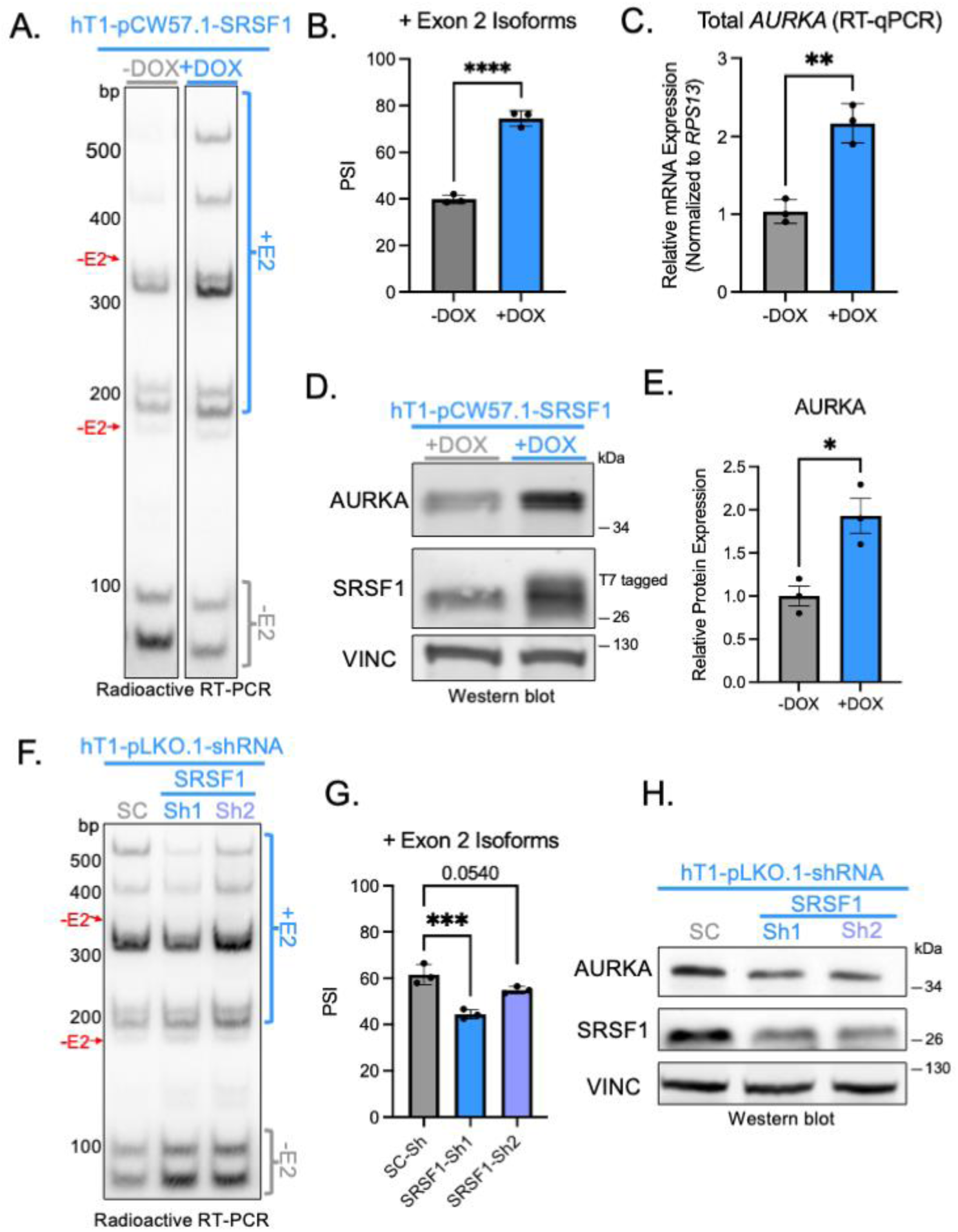
SRSF1 regulates human *AURKA* +E2 isoforms in PDAC organoids (A) Radioactive RT-PCR of *AURKA* 5’UTR from hT1-pCW57.1-SRSF1 ± 2 μg/ml DOX for 48 h. Lanes were cropped from the same gel. (B) PSI quantification of +E2 isoforms from (A) (n = 3 biological replicates). (C) RT-qPCR of total *AURKA* from (A) normalized to *RPS13* (n = 3 biological replicates). (D) Western blot of AURKA, SRSF1, and VINC from hT1-pCW57.1-SRSF1 ±DOX treatment as in (A). (E) Relative protein expression of AURKA compared to -DOX normalized to VINC (n = 3 biological replicates). (F) Radioactive RT-PCR of *AURKA* 5’UTR from hT1-pLKO.1-shRNA transduced with either SC-shRNA, SRSF1-sh1, or SRSF1-sh2. SC, scrambled control; sh, short-hairpin RNA. (G) PSI quantification of +E2 isoforms from (F) (n = 3 biological replicates). (H) Western blotting of AURKA, SRSF1, and VINC from hT1-pLKO.1-shRNA organoids as in (F). Data shown as mean ± SD. *p ≤ 0.05, **p ≤ 0.01, ***p ≤ 0.001 ****p ≤ 0.0001 by Student’s t test (B,C, and E) or one-way ANOVA (G). See also Table S1 and Figure S4.

Knockdown of SRSF1 using shRNA transduced into hT1 organoids resulted in a shift from +E2 isoforms to -E2 isoforms, and decreased AURKA protein levels, though these effects were less pronounced than in the overexpression experiments, likely reflecting SR-protein functional redundancy (Figure 3F-H).^52^

*SRSF1* and *AURKA* mRNA levels were correlated in PDAC tumors (TCGA) and pancreatic cancer cell lines (DepMap Portal) (Figure S4C-D). This correlation was also evident in our organoid models (Pearson coefficient = 0.9318) (Figure S4E), reinforcing the regulatory link between SRSF1 and *AURKA* splicing.

### Alternative splicing within the *AURKA* 5’UTR increases protein expression through a splicing-dependent mechanism

We next sought to determine whether the increase in AURKA protein expression primarily reflects higher total mRNA levels or whether the +E2 isoforms are translated more efficiently. One group reported that some +E2 isoforms (NM_198436.3 and NM_198434.3; See Figure 2B) are more translationally active in colorectal cancer through EGFR signaling,^37^ but later also reported that hnRNP Q1, which is upregulated in colorectal cancer and mediates tumorigenesis, translationally activates all the observable *AURKA* 5’UTR isoforms in their model, especially the shorter -E2 isoforms (NM_198437.3 and NM_198435.3).^53^

To test the translation potential of *AURKA* 5’UTR isoforms in the PDAC context, we generated luciferase reporters comprising cDNAs for the five major *AURKA* 5’UTR isoforms expressed in PDAC organoids (Figure S5A). We transfected these reporters into the PDAC 2D cell line SUIT2, which harbors the *KRAS^G12D^* mutation and is easier to handle than organoids. All the isoforms exhibited similar Firefly activity, indicating no intrinsic sequence-driven translational advantage for the +E2 isoforms in PDAC, at least when expressed from cDNAs (Figure S5B). We also found no differences in endogenous mRNA isoform half-lives, following transcription inhibition with actinomycin D, regardless of exon 2 inclusion status (Figure S5C-D). However, polysome fractionation revealed that the endogenous +E2 isoforms associated with heavier polysomes, suggesting higher translational rates (Figure S5E-H).

The observation that inclusion of exon 2 correlates with higher mRNA and protein led us to investigate potential splicing-dependent mechanisms of gene regulation. Splicing-mediated removal of introns, particularly promoter-proximal introns, enhances gene expression,^54–58^ and can positively affect all stages of mRNA processing, including transcription, stability, export, and translation.^59–62^ Efficiently spliced introns in the 5’UTR (tested in the context of single-intron reporters) generally enhance gene expression independently of intron sequence, with the primary result being increased mRNA levels.^59^ This general phenomenon of splicing-mediated removal of introns stimulating gene expression—whether or not in the 5’UTR—is referred to as intron-mediated enhancement (IME).^59,63^ A related phenomenon termed exon-mediated activation of transcription starts (EMATS), which involves splicing of evolutionarily recent alternative exons near a promoter, often within the 5’UTR, also stimulates gene expression.^64,65^

Given that exons 2 and 3 of *AURKA* are evolutionarily recent, reside in the 5’UTR, and SRSF1 activity correlates with exon 2 inclusion and increased total *AURKA* mRNA, we hypothesized that inclusion of exon 2 within the *AURKA* 5’UTR stimulates gene expression. As we noted above (Figure S3C), the relatively weak exon 2 3’ss remains unmutated from the ancestral AluJo sequence, suggesting a potentially important regulatory region. To test this hypothesis, we generated splicing-competent minigene reporters by inserting the entire *AURKA* 5’UTR between the *PGK* promoter and the Firefly start codon (Figure S6A). Strengthening the exon 2 3’ss (MAXENT 2.41 → 10.05), resulted in greater exon 2 inclusion (average +22% ΔPSI), leading to enhanced *Firefly* mRNA expression (average 2.9-fold) and a 2.5-fold increase in Firefly activity (Figure 4A-E).

**Figure 4.**
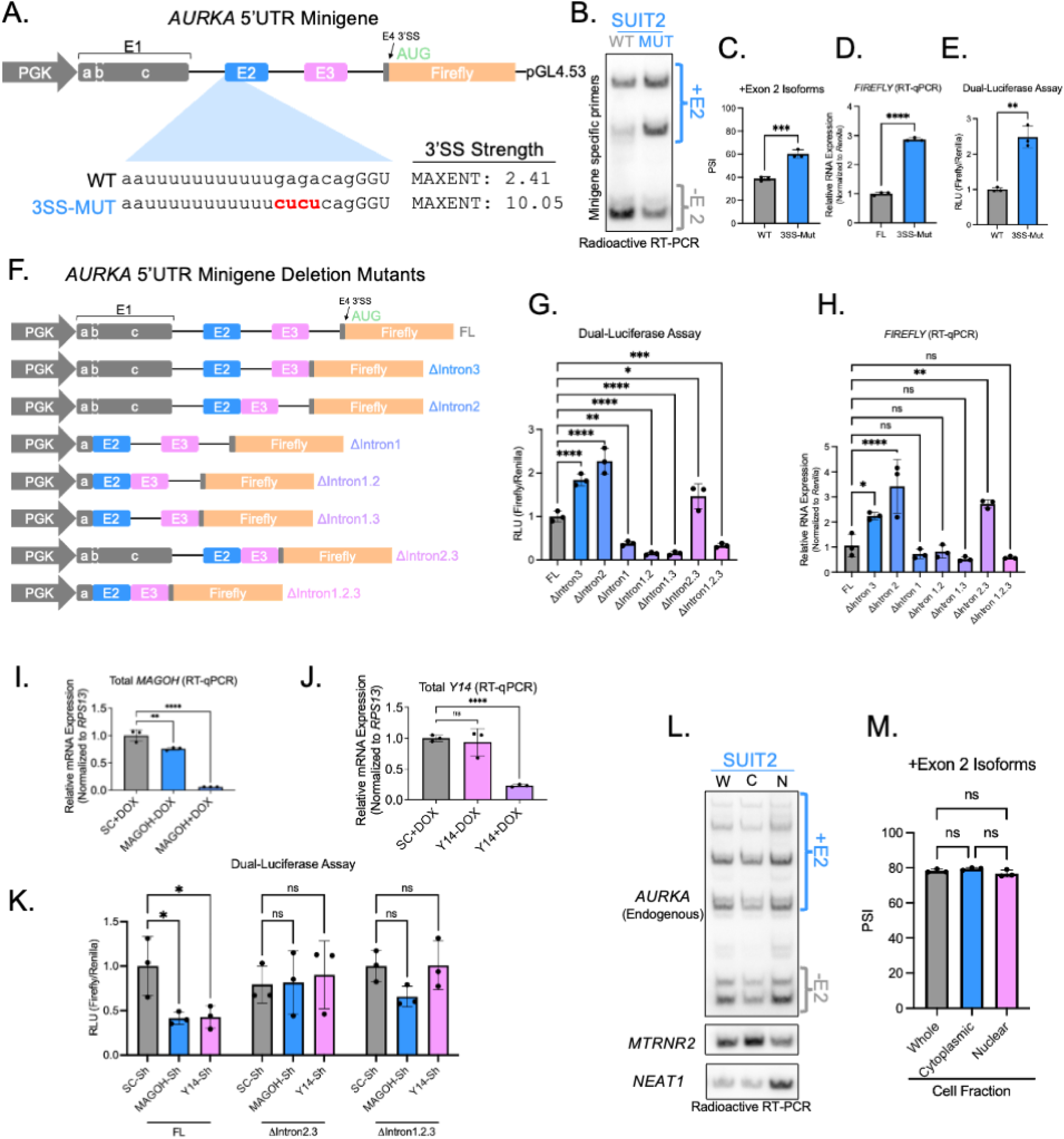
Alternative splicing within the *AURKA* 5’UTR increases protein expression through a splicing-dependent mechanism (A) Schematic of *AURKA* 5’UTR minigene (top). Wild-type (WT) and 3’ss mutant (3SS-Mut) sequences, (bottom). Maxent scores calculated using MaxEntScan::score3ss. (B) Radioactive RT-PCR of minigenes transfected in SUIT2 cells (24 h). (C) PSI quantification of +E2 isoforms from (B) (n = 3 biological replicates). (D) RT-qPCR of total *Firefly* from (B) normalized to *Renilla* (n = 3 biological replicates). (E) Dual-luciferase assay from SUIT2 cells transfected as in (B). Firefly activity normalized to Renilla (n = 3 biological replicates). RLU, Relative Luciferase Units. (F) Schematic of minigene deletion mutants. FL, full length. (G-H) Dual-luciferase assay (H) and RT-qPCR (H) from SUIT2 cells transfected with mutants from (F). Firefly/*Firefly* normalized to Renilla/*Renilla* (n = 3 biological replicates). (I-J) RT-qPCR of *MAGOH* (H) and *Y14* (I) from SUIT2-Tet-pLKO.1-shRNA after ±2 μg/ml DOX treatment for 48 h. (K) Dual-luciferase assay from SUIT2-Tet-pLKO-shRNA cell lines +2 μg/ml DOX for 48 h and transfection of FL, ΔIntron 2.3, or ΔIntron 1.2.3 for 24 h (from (F)). (L) Radioactive RT-PCR of *AURKA* 5’UTR, *MTRNR2*, and *NEAT1* from SUIT2 cells after cellular fractionation. W, whole-cell fraction; C, cytoplasmic fraction; N, nuclear fraction. (M) PSI quantification of +E2 isoforms from (L). Data shown as mean ± SD. ns = non-significant, *p ≤ 0.05, **p ≤ 0.01, ***p ≤ 0.001 ****p ≤ 0.0001 by Student’s t test (C-E) or one-way ANOVA (G-K and N). See also Figure S5-7.

To evaluate the contribution of 5’UTR introns to *AURKA* expression, we sequentially deleted individual introns, generated double-deletion mutants, and created an intronless construct (Figure 4F). Deletion of intron 3 resulted in a 1.8-fold increase in Firefly activity and a 2.2-fold increase in *Firefly* mRNA levels, compared to the full-length (FL) construct (Figure 4G-H). Deletion of intron 2 further stimulated reporter expression, yielding a 2.3-fold increase in Firefly activity and 3.4-fold increase in *Firefly* mRNA expression (Figure 4G-H). Strikingly, deletion of intron 1 resulted in a pronounced reduction in Firefly activity across all single, double, or triple intron-deletion mutants, with an average decrease of 76% (Figure 4G). This reduction in activity was accompanied by a modest though not significant average reduction of 34% in mRNA levels across the intron 1 deletion mutants, suggesting that splicing of intron 1 is critical and has potent IME effects on expression (Figure 4H). The ΔIntron 2.3 mutant displayed similar expression patterns as the ΔIntron 3 or ΔIntron 2 mutants, with a 1.5-fold increase in Firefly activity and a 2.7-fold increase in mRNA levels compared to FL (Figure 4G-H).

Splicing analysis confirmed that all constructs expressed functional splice variants, with no evidence of intron retention (Figure S6B), which could otherwise introduce intronic upstream open reading frames (uORFs). Interestingly, the ΔIntron 2 mutant resulted in complete skipping of exon 2, generating the shortest 5’UTR isoform (ISO1, see Figure S5A). This mutant exhibited the highest mRNA and protein levels, which seems paradoxical, given that the +E2 isoforms are upregulated in PDAC, correlate with higher mRNA levels upon SRSF1 overexpression, and associate with heavier polysomes. However, in this context, ISO1 appears to be the most efficiently expressed *AURKA* isoform.

To address this paradoxical observation, we analyzed the predicted RNA secondary structures of each deletion mutant (Figure S7A-G). Deletion of introns 2 or 3 markedly reduced the predicted dsRNA structure formed by the AluSq2 and AluSp elements, whereas deletion of intron 1 largely preserved it. This dsRNA structure may hinder efficient *AURKA* splicing, accounting for the mRNA accumulation when the structure is abolished. Hence, ΔIntron 2 expresses the optimal isoform (ISO1), with the highest expression levels reflecting improved splicing efficiency (See Discussion).

Splicing can also stimulate translation through the deposition of the exon-junction complex (EJC) and subsequent assembly into polysomes.^61^ We tested the EJC’s role in *AURKA* translation by knockdown of MAGOH or Y14, two core EJC components (Figure 4I-J).^66^ EJC depletion reduced FL reporter expression by >50%, whereas ΔIntron 2.3—which should have only one EJC—and the intronless construct remained unaffected, indicating that either cumulative or site-specific EJC deposition along the 5’UTR enhances protein expression (Figure 4K). Nuclear/cytoplasmic fractionation showed no difference in endogenous or minigene *AURKA* mRNA isoform subcellular localization, suggesting that the 5’UTR EJC stimulates translation without affecting mRNA export (Figure 4L-M, S6C).

Taken together, these findings demonstrate that *AURKA* 5’UTR alternative splicing enhances protein expression by: i) increasing mRNA levels via efficient splicing; and ii) enhancing translation through the deposition of EJCs at particular locations. These effects support a splicing-dependent mechanism, explaining why the pre-spliced +E2 cDNA reporters showed no translation advantage.

### The SRSF1-AURKA-MYC circuit is independent of AURKA kinase activity and dependent on AURKA protein levels

We next investigated the role of the SRSF1-AURKA-MYC circuit in PDAC. AURKA knockdown was previously shown to reduce SRSF1 protein levels in HeLa cells post-translationally via AURKA kinase activity.^34^ Additionally, AURKA can phosphorylate SR proteins, including SRSF1, and influence AS networks.^67^ Furthermore, AURKA stabilizes MYC and N-MYC in the nucleus by preventing their proteasomal degradation,^27,30,31,68^, whereas MYC transcriptionally activates both *AURKA* and *SRSF1*, establishing a multilayered genetic circuit.^6,29,32,69,70^ Knockdown of either SRSF1 or AURKA led to a substantial reduction in the other one, as well as in MYC protein levels, confirming that these oncogenic factors are tightly linked at the protein level in PDAC (Figure 5A-B).

**Figure 5.**
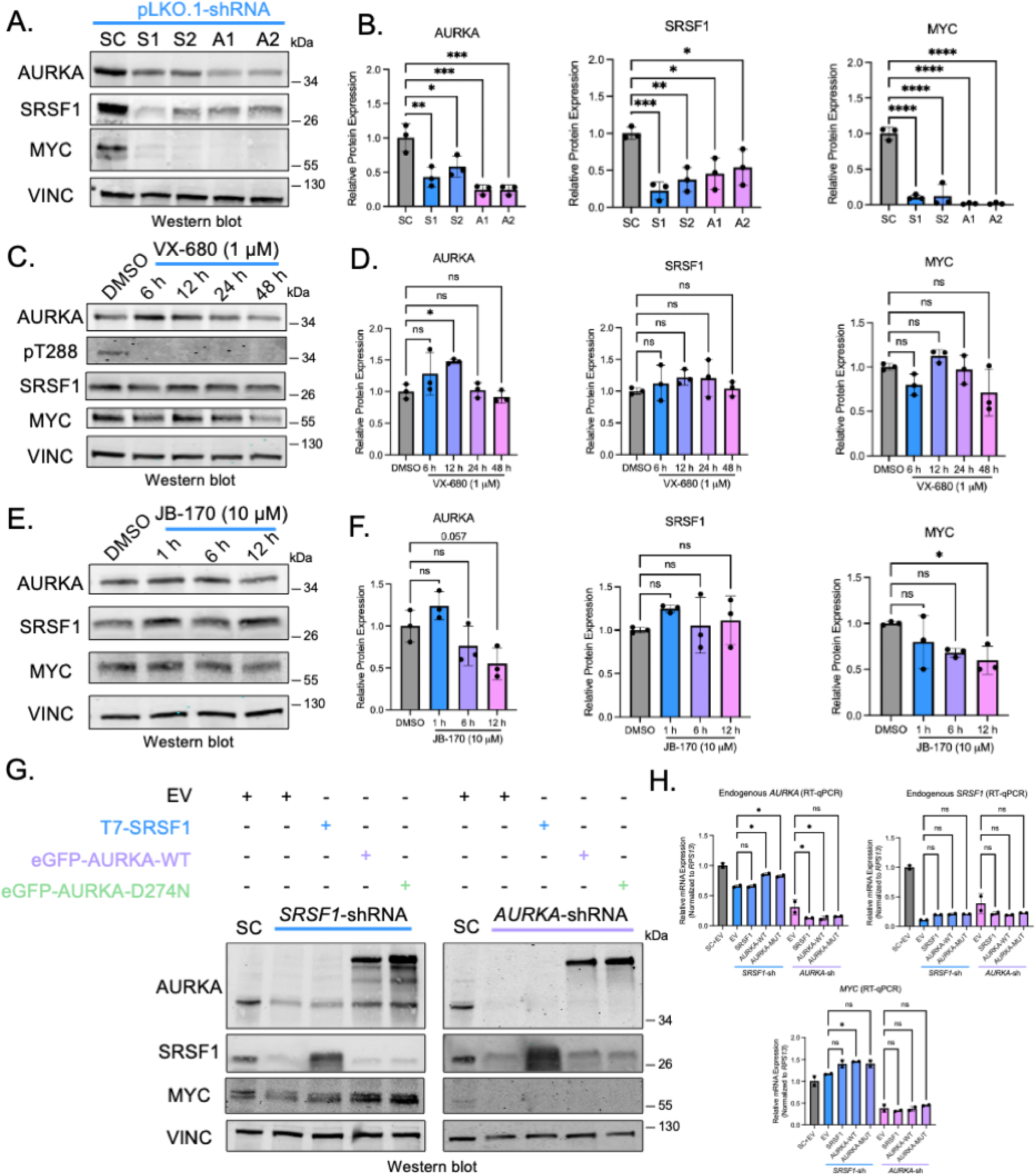
The SRSF1-AURKA-MYC circuit is independent of AURKA kinase activity and dependent on AURKA protein levels (A) Western blotting of AURKA, SRSF1, MYC, and VINC from SUIT2-pLKO.2-shRNA cell after lentiviral transduction of shRNA targeting *SRSF1* (S1 and S2) or *AURKA* (A1 and A2), or a scrambled-sequence control shRNA (SC). (B) Relative protein expression compared to SC for AURKA (left), SRSF1 (middle), and MYC (right) from (A) normalized to VINC (n = 3 biological replicates). (C) Western blotting of AURKA, AURKA phosphorylated at Thr288 (pT288), SRSF1, MYC, and VINC from SUIT2 cells after treatment with VX-680 at 1 μM or DMSO. (D) Relative protein expression compared to DMSO for AURKA (left), SRSF1 (middle), and MYC (right) from (C) normalized to VINC (n = 3 biological replicates). (E) Western blotting of AURKA, SRSF1, MYC, and VINC from SUIT2 cells after treatment with AURKA PROTAC JB-170 at 10 μM. (F) Relative protein expression compared to DMSO for AURKA (left), SRSF1 (middle), and MYC (right) from (E) normalized to VINC (n = 3 biological replicates). (G) Western blotting of AURKA, SRSF1, MYC, and VINC from SUIT2 cells transduced with lentiviral shRNAs targeting *SRSF1, AURKA*, or SC shRNA for 72 h. After 72 h, cells were transfected with shRNA-resistant plasmids: empty vector (EV), T7-SRSF1, eGFP-AURKA-WT, or eGFP-AURKA-D274N (catalytically dead mutant). (H) RT-qPCR of endogenous total mRNA for *AURKA* (left), *SRSF1* (right), and *MYC* (bottom) from SUIT2 cells as in (G). Normalized to *RPS13* (n = 2 biological replicates). Data shown as mean ± SD. ns = non-significant, *p ≤ 0.05, **p ≤ 0.01, ***p ≤ 0.001 ****p ≤ 0.0001 one-way ANOVA (B, D, F, H). See also Figure S8.

To determine whether AURKA’s kinase activity is required for this circuit, we treated PDAC cells with the AURKA inhibitor VX-680^71^ (1 μM) and performed a time-course analysis. Despite complete inhibition of T288 phosphorylation (an autophosphorylation marker of AURKA kinase activity^72,73^), SRSF1 and MYC remained unchanged, indicating that AURKA’s oncogenic role in this circuit is kinase-independent (Figure 5C-D). However, at higher VX-680 concentrations and extended treatment times, we did observe downregulation of SRSF1 and MYC. This effect, however, coincided with the complete loss of AURKA protein, suggesting that SRSF1 and MYC expression depend on AURKA protein levels, rather than its kinase activity (Figure S8A-B).

Next, we used the AURKA-specific PROTAC degrader JB170 to induce acute AURKA depletion.^68^ After 12 h, AURKA levels decreased by 50%, leading to a proportional decrease in MYC, whereas SRSF1 remained initially unaffected (Figure 5E-F). This observation supports the model that AURKA stabilizes MYC by masking its phospho-T58 degradation signal, such that its depletion exposes MYC to proteasomal degradation^30,74^. The delayed downregulation of SRSF1 reflects its dependence on MYC-driven transcription, requiring additional time for *SRSF1* mRNA and/or SRSF1 protein turnover.

To refine this model, we performed rescue experiments with shRNA-resistant *SRSF1* and *AURKA* cDNA constructs, 72 h after lentiviral shRNA knockdown. Both WT and kinase-dead (D274N^27^) AURKA mutants restored MYC levels in SRSF1-knockdown cells, confirming that AURKA’s regulatory role is independent of its kinase activity (Figure 5G and H). However, in AURKA-knockdown cells, no condition restored protein expression, suggesting that after extensive depletion of AURKA and MYC, the AURKA constructs are not sufficient to revive the circuit at this time point.

These findings establish that the SRSF1-AURKA-MYC feedback loop is dependent on AURKA protein expression, rather than its kinase activity.

### Uniformly modified splice-switching ASO targeting exon 2 induces its skipping and reduces total *AURKA* mRNA levels

We next investigated whether targeting *AURKA* 5’UTR splicing and impeding splicing-dependent mRNA expression could effectively downregulate AURKA protein levels and disrupt the SRSF1-AURKA-MYC oncogenic circuit. We designed splice-switching ASOs to induce exon 2 skipping. We used uniformly modified 2’-O-methoxyethyl (MOE) ASOs with a phosphorothioate (PS) backbone and 5-methylcytosine (5mC) modifications. We tested six ASOs targeting either the 3’ss or the 5’ss of exon 2 (Figure 6A). Transfection at 100 nM for 48 h into SUIT2 PDAC cells, followed by radioactive RT-PCR, revealed that ASO-3 and ASO-6 elicited the most exon 2 skipping, albeit modestly (Figure 6B-C). Surprisingly, despite their uniform design, which does not elicit RNase-H cleavage of the target RNA, all the ASOs reduced the total *AURKA* mRNA levels, with ASO-3 almost completely eliminating *AURKA* expression (Figure 6D). These ASOs also reduced cell viability, with ASO-3 exhibiting the strongest effects (Figure 6E).

**Figure 6.**
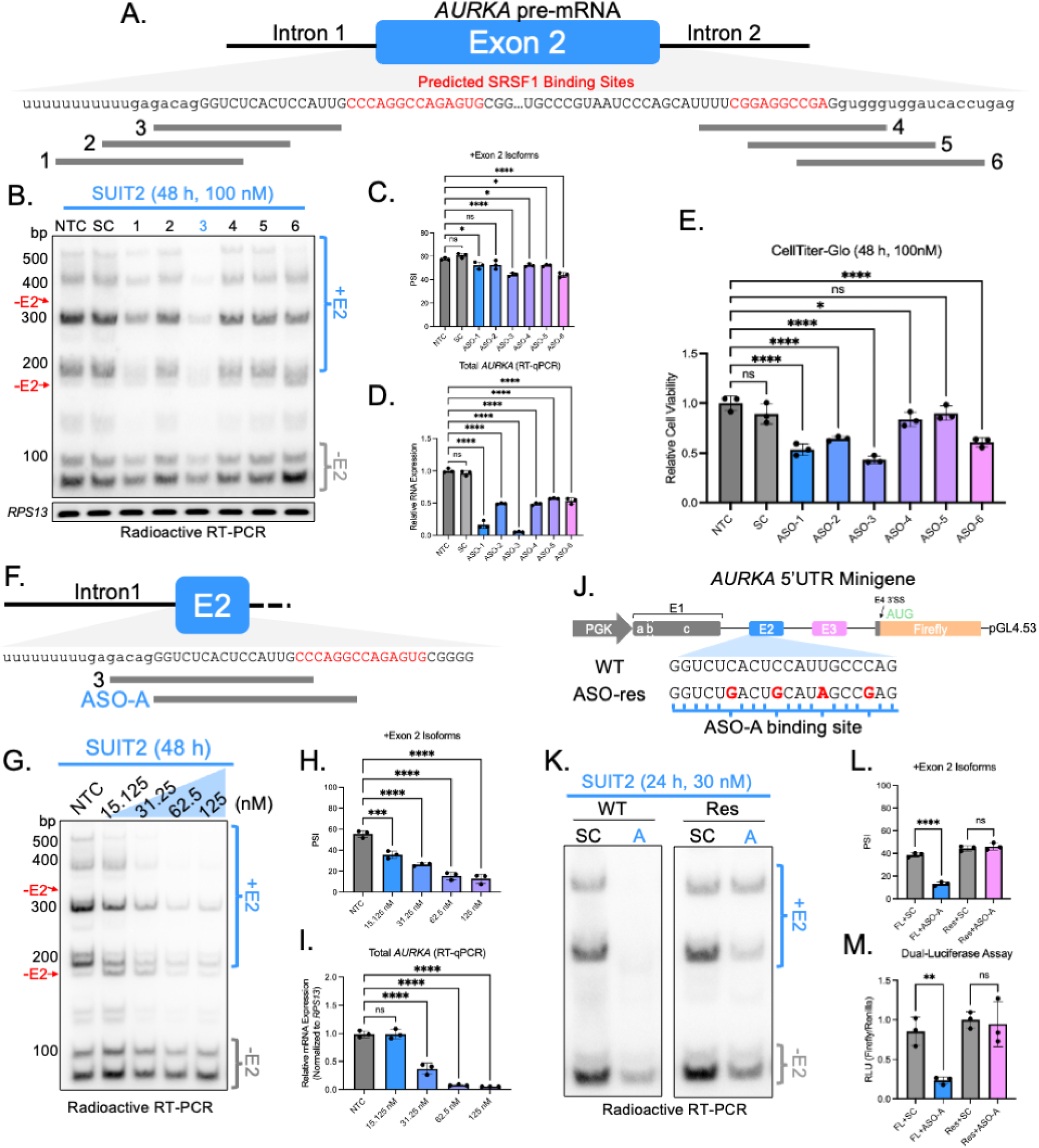
Uniformly modified splice-switching ASO targeting exon 2 induces its skipping and reduces total *AURKA* mRNA levels (A) Schematic of exon 2 ASO binding sites. SRSF1 ESEs (ESEfinder 3.0) in red. (B) Radioactive RT-PCR of *AURKA* 5’UTR from SUIT2 cells after transfection of ASOs at 100 nM for 48 h (top). RT-PCR of *RPS13* (bottom). No-treatment control (NTC, Lipofectamine 3000 only); SC, scrambled control; 1, ASO-1; 2, ASO-2; 3, ASO-3; 4, ASO-4; 5, ASO-5; 6, ASO-6. (C and D) PSI quantification of +E2 isoforms (C) and RT-qPCR of total *AURKA* normalized to *RPS13* (D) from (B) (n = 3 biological replicates). (E) Cell-viability assay of SUIT2 cells transfected with ASOs as in (B) normalized to day 0 reading (n = 3 biological replicates). (F) Schematic of ASO-A (lead ASO) and ASO-3 in exon 2. (G) Radioactive RT-PCR of *AURKA* 5’UTR from SUIT2 cells after transfection of ASO-A. (H and I) PSI quantification of +E2 isoforms (H) and RT-qPCR of total *AURKA* normalized to *RPS13* from (G) (n = 3 biological replicates). (J) Schematic of *AURKA* 5’UTR minigene. WT and ASO-resistant (ASO-res) sequences are shown, with the mutated positions indicated in red, corresponding to ASO mismatches. (K) Radioactive RT-PCR of *AURKA* minigene from SUIT2 cells co-transfected with minigenes from (J) and ASOs at 30 nM for 24 h. A, ASO-A. (L and M) PSI quantification of +E2 isoforms (L) and dual-luciferase assay (M) of (K) (n = 3 biological replicates). Data shown as mean ± SD. ns = non-significant, *p ≤ 0.05, **p ≤ 0.01, ***p ≤ 0.001 ****p ≤ 0.0001 one-way ANOVA (C, D, E, H, I, L, M). See also Figure S9.

To optimize exon 2 skipping, we designed three additional ASOs that overlap ASO-3 (Figure S9A) and target a conserved SRSF1 ESE motif (Figure S1C), a region likely important for *AURKA* splicing regulation. All three ASOs effectively induced exon 2 skipping, reduced total *AURKA* mRNA, depleted AURKA, SRSF1, and MYC protein levels, and significantly reduced cell viability, with ASO-3.1 showing the strongest effects (Figure S9B-F). Based on these findings, we selected ASO-3.1 (renamed ASO-A) as our lead ASO.

To assess ASO-A’s specificity, we performed a dose-response experiment, which confirmed dose-dependent splice switching and total *AURKA* mRNA depletion (Figure 6F-I). A time-course analysis further demonstrated that ASO-A downregulates total *AURKA* mRNA, along with AURKA, SRSF1, and MYC protein levels, as early as 24 h, with maximal effects observed at 48 h (Figure S9G-J).

To further validate ASO-A’s specificity, we introduced four nucleotide substitutions in the ASO- complementary region of our *AURKA* minigene reporter, generating an ASO-resistant mutant. The resistant construct remained unaffected by ASO-A-mediated splice switching and showed no reduction in luciferase signal, confirming that ASO-A specifically targets *AURKA* (Figure 6J-M).

To ensure that exon 2 skipping directly leads to AURKA protein depletion, rather than depletion being the result of off-target effects, we tested ASO-6, which binds the 5’ss of exon 2 but induces less total *AURKA* mRNA knockdown. In SUIT2 cells, ASO-6 efficiently induced exon 2 skipping and led to AURKA, SRSF1, and MYC protein depletion, despite having no effect on total *AURKA* mRNA levels when transfected at 65 nM. However, ASO-A remained significantly more potent, even at 30 nM (Figure S9K-N).

### ASO-A collapses the SRSF1-AURKA-MYC circuit, induces apoptosis, and overcomes KRAS^G12D^ inhibitor resistance in PDAC

Using the optimal 65 nM dose, which effectively induced *AURKA* exon 2 skipping and reduced total *AURKA* mRNA, we assessed the impact of ASO-A on the SRSF1-AURKA-MYC circuit. Transfection of SUIT2 cells for 48 h with ASO-A resulted in extensive *AURKA* mRNA depletion (Figure 7A-C), with modest reductions in *SRSF1* and *MYC* mRNA levels (Figure 7D-E). At the protein level, ASO-A treatment led to the nearly complete loss of AURKA, SRSF1, and MYC (Figure 7F), correlating with significant reductions in cell viability and increased apoptosis (Figure 7G-H). We also validated these findings in hT1 organoids, in which ASO-A similarly induced exon 2 skipping, disrupted the SRSF1-AURKA-MYC circuit, and reduced viability (Figure S10A-G).

**Figure 7.**
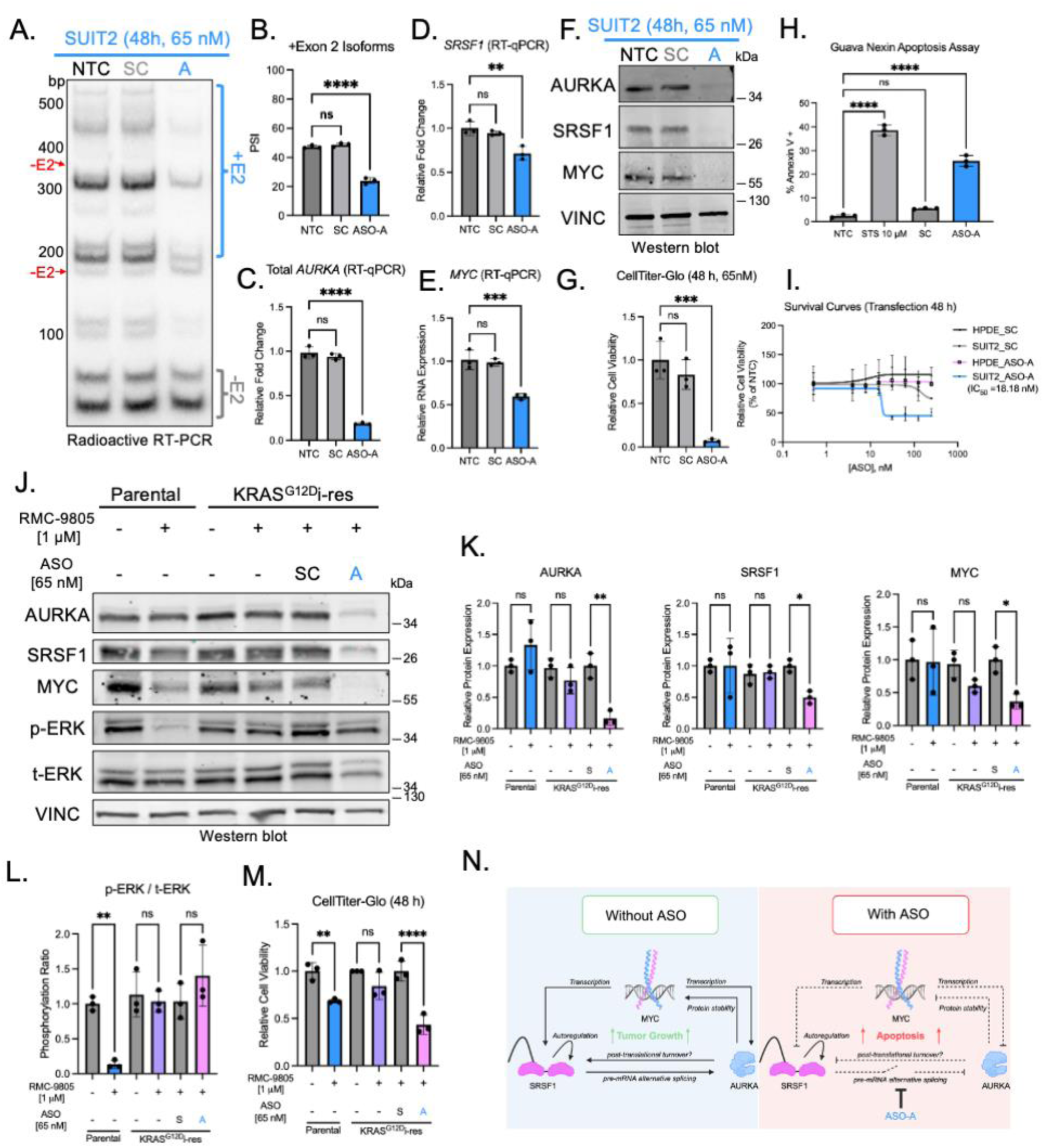
ASO-A collapses the SRSF1-AURKA-MYC circuit, induces apoptosis, and overcomes KRAS^G12D^ inhibitor resistance in PDAC (A) Radioactive RT-PCR of *AURKA* 5’UTR from SUIT2 cells after ASO transfection. NTC, non-treated control (Lipofectamine 3000 only); SC, scrambled control; A, ASO-A. (B-E) PSI quantification of +E2 isoforms (B) and RT-qPCR of total *AURKA* (C), *SRSF1* (D), and *MYC* (E) normalized to *RPS13* from (A) (n = 3 biological replicates). (F) Western blotting of AURKA, SRSF1, MYC, and VINC after ASO treatment as in (A). (G) Cell-viability assay of SUIT2 cells after ASO treatment as in (A), normalized to day 0 (n = 3 biological replicates). (H) Apoptosis assay of SUIT2 cells after ASO treatment as in (A). Staurosporine (STS) at 10 μM for 24 h as a positive control (n = 3 biological replicates). (I) Dose-response in SUIT2 vs. HPDE cells. Points are the mean ± SD (n = 3 biological replicates). (J) Western blotting of AURKA, SRSF1, MYC, phosphorylated and total ERK1/2 (p-ERK and t-ERK, respectively), and VINC from SUIT2 parental cells and SUIT2 RMC-9805-resistant cells. Cells were either treated with DMSO, RMC-9805 [1 μM], or a combination of RMC-9805 [1 μM] and SC or ASO-A [65 nM] for 48 h. (K) Relative protein expression compared to each indicated control, for AURKA (left), SRSF1 (middle), and MYC (right) from (J) normalized to VINC (n = 3 biological replicates). (L) Quantification of the ratio of p-ERK/t-ERK from (J) normalized to VINC; p-ERK was then normalized to t-ERK, and the ratio was then normalized to each indicated control (n = 3 biological replicates). (M) Cell-viability assay from SUIT2-parental and SUIT2-RMC-9805-resistant cells treated as in (I) normalized to day 0 (n = 3 biological replicates). (N) Schematic of SRSF1-AURKA-MYC circuit. Data shown as mean ± SD. ns = non-significant, *p ≤ 0.05, **p ≤ 0.01, ***p ≤ 0.001 ****p ≤ 0.0001 one-way ANOVA (B, C, D, E, G, H, K, L, M). See also Figures S10 and S11.

Given that the -E2 isoforms are the predominant *AURKA* transcripts in normal tissues besides testes (Figure S1D), we next assessed whether ASO-A selectively targets PDAC cells over normal cells. Dose-response analysis revealed that SUIT2 cells were sensitive to ASO-A, with an IC50 of 18 nM, whereas the immortalized human pancreatic ductal cell line HPDE was unaffected (Figure 7I). To rule out differences in transfection efficiency, we transfected both lines with Nusinersen, an ASO that promotes *SMN2* exon 7 inclusion,^75^ and observed nearly complete exon 7 inclusion in both lines, indicating equivalent delivery (Figure S10H).

Taking into account that AURKA and MYC mediate KRAS-inhibitor resistance,^14,21,76^ we next tested whether ASO-A is effective in KRAS inhibitor-resistant cells. Using the KRAS^G12D^ inhibitor RMC-9805^77^, we first generated drug-resistant cells by incubating SUIT2 cells with RMC-9805 at 10 μM for one month. We then confirmed that the parental SUIT2 cells responded to treatment, showing suppression of ERK1/2 phosphorylation (pERK) and reduced cell viability, whereas the resistant cells maintained MAPK signaling and showed no difference in cell viability (Figure 7J-M). AURKA, SRSF1, and MYC protein levels remained static between parental and resistant cell lines, regardless of inhibitor presence. In contrast, ASO-A treatment effectively depleted AURKA, SRSF1, and MYC proteins in the drug-resistant cells, this time with no impact on ERK1/2 phosphorylation, yet it still resulted in robust cell-viability defects (Figure 7J-M). These results demonstrate that ASO-A mediated depletion of *AURKA* collapses the SRSF1-AURKA-MYC circuit independently from the MAPK pathway.

Considering that *AURKA* overexpression is linked to poor survival in kidney and liver cancers (Figure S11A- B), we further tested ASO-A in two renal cell carcinoma (RCC) lines and two hepatocellular carcinoma (HCC) lines, as well as in three additional PDAC cell lines. All these cell lines expressed *AURKA* +E2 isoforms, and ASO-A effectively induced exon 2 skipping, reduced AURKA protein expression, and reduced cell viability across all the lines, suggesting that this strategy could be applicable beyond PDAC (Figure S11C-F). Interestingly, SRSF1 and MYC protein levels were less influenced by AURKA in RCC and HCC cell lines than in PDAC cell lines, suggesting that the SRSF1-AURKA-MYC circuit may be context-specific.

In conclusion, ASO-A-mediated modulation of *AURKA* 5’UTR AS collapses the SRSF1-AURKA-MYC circuit and induces apoptosis (Figure 7N). These results suggest that targeting *AURKA* splicing could overcome KRAS-inhibitor resistance and highlight a potential combination therapy.

## DISCUSSION

We uncovered a previously unrecognized oncogenic circuit involving SRSF1, AURKA, and MYC, and demonstrated that *AURKA* 5′UTR AS is a critical regulatory step in AURKA protein expression. Specifically, we showed that SRSF1 promotes exon 2 inclusion in the *AURKA* 5′UTR, enhancing mRNA accumulation and EJC deposition to promote translation via IME- and EMATS-related mechanisms. Our findings highlight a fundamental splicing-dependent mechanism that regulates *AURKA* expression and suggest how this process is buffered by Alu-derived dsRNA structures embedded within the *AURKA* 5′UTR.

We described how human *AURKA* exons 2 and 3 originated from exonized Alu elements and potentially form extensive dsRNA structure in the pre-mRNA that appears to act as a natural brake on *AURKA* expression by impeding efficient splicing. We found that efficient inclusion of exon 2, either through SRSF1 activity or by strengthening the 3’ss, increased *AURKA* mRNA and AURKA protein expression, which is consistent with EMATS, suggesting an exon-centric regulatory mechanism of *AURKA* gene expression.

However, deletion of intron 2—which spans most of AluSp—resulted in enhanced mRNA and protein expression, with the predominant transcript being the shortest isoforms (NM_198437.3 and NM_198435.3). Thus, in the absence of Alu-derived dsRNA, splicing preferentially and efficiently generates the shortest isoforms, resulting in higher expression, which is more representative of an IME-type regulation. Under physiological conditions, the generation of this splice isoform is constrained by the inhibitory dsRNA structure. Upregulated SRSF1 facilitates efficient exon 2 splicing within the 5’UTR, enabling splicing-dependent mRNA accumulation and EJC deposition to enhance AURKA protein expression, utilizing both IME and EMATS mechanisms. Since the CDS and 3’UTR are constitutively spliced,^32^ only EJC-deposition differences in the 5’UTR are expected to influence translation.

Beyond splicing, SRSF1 promotes mTORC1 signaling by stimulating S6K and 4E-BP1 phosphorylation.^78,79^ Through this mechanism, SRSF1 promotes translation, both indirectly via 4E-BP1 inactivation, and directly by remaining bound to cytoplasmic mRNA’s.^79,80^ A previous study used exon-junction microarrays and showed that SRSF1 promotes exon 2 inclusion in *AURKA*, although *AURKA* was not directly translationally activated by SRSF1.^81^ Our findings suggest that SRSF1 primarily regulates *AURKA* by promoting exon 2 inclusion, thereby increasing mRNA accumulation and EJC deposition.

This mechanism of increased AURKA protein expression downstream of SRSF1 likely differs in mouse *Aurka*, as we observed elevated protein levels without a corresponding increase in mRNA. This difference may be attributed to EJC deposition, direct translational control by SRSF1, or enhanced mRNA stability. Despite these species-specific differences, it is striking that in both human and mouse cancer models, SRSF1 converges on *AURKA/Aurka* regulation through AS in the 5’UTR—even in the absence of strict sequence conservation and the presence of the Alu-elements in the human 5’UTR.

Other mechanisms we did not directly test include: IRES-dependent translation, potentially facilitated by Alu-derived RNA structures in the 5’UTR^82^—though both *AURKA* -E2 (NM_198437.3 and NM_198435.3) and +E2 (NM_198436.3 and NM_198434.3) isoforms exhibit IRES activity^53^; microRNAs that might bind to structured regions in the 5’UTR and enhance stability^83^; and differences in translation initiation rates linked to 5’UTR length and structure.^84^ 5’UTR variants generated by AS can also influence mRNA stability^85^, as we previously showed for *IL1R1,* another SRSF1 target.^6^ However, in the case of *AURKA*, we did not detect significant differences in mRNA half-life among isoforms.

We established a splicing-dependent link between SRSF1, AURKA, and MYC, defining an oncogenic feedback loop. Whereas it was previously shown that AURKA positively regulates both SRSF1 and MYC,^28,30,31,34^ and that MYC transcriptionally activates *SRSF1* and *AURKA*,^6,28,29,69,70^ our findings here demonstrated that SRSF1 directly regulates *AURKA* expression via 5’UTR alternative splicing, thereby closing the loop on this oncogenic circuit. Our findings provide additional insight into why AURKA kinase inhibitors have had limited clinical efficacy, as we found that this circuit is independent of AURKA’s kinase activity and highly dependent on AURKA protein levels. These findings underscore the importance of targeting AURKA’s non-catalytic functions as a therapeutic strategy.

Given the preferential expression of the +E2 *AURKA* isoforms in PDAC, we used splice-switching ASOs to target this tumor-specific alternative-splicing event. Several ASOs induced exon 2 skipping, and, surprisingly, reduced *AURKA* mRNA, despite their uniform design, which does not elicit RNase H cleavage. This result strongly supports our model that *AURKA* 5′UTR exon 2 inclusion enhances AURKA protein expression by promoting mRNA accumulation and translation. The exon 2 3’ss region proved highly responsive to ASOs, likely due to strong SRSF1 motifs and greater accessibility than the 5’ss, which is masked by Alu-derived dsRNA. ASO-6, targeting the 5’ss, reduced AURKA protein but not mRNA levels— potentially by disrupting dsRNA structure and relieving splicing repression—allowing mRNA accumulation but generating suboptimally translated isoforms, resulting in reduced protein output.

Beyond PDAC, we showed that high *AURKA* mRNA expression correlates with poor survival in HCC and RCC, and that ASO-A is effective in these cancer types as well. As MYC and AURKA are implicated in chemoresistance and KRAS-inhibitor resistance,^19,21,27,32,76,86–89^ targeting *AURKA* exon 2 splicing could broadly counter tumor plasticity and resistance mechanisms. Notably, KRAS^G12D^ inhibitor-resistant cells remained sensitive to ASO-A-mediated collapse of the SRSF1-AURKA-MYC circuit, underscoring the therapeutic relevance of this approach. Although siRNAs have been used to target *AURKA* exon 2 in colorectal cancer,^36^ this is the first application of a splicing-directed approach using uniformly modified ASOs to abrogate AURKA expression. By disrupting *AURKA* 5′UTR splicing, we leveraged the presence of Alu dsRNA to interfere with splicing-mediated mRNA expression, establishing a therapeutic strategy for suppressing *AURKA* expression.

We believe that splice-switching ASOs, in this context, offer a distinct therapeutic advantage over other oligonucleotide modalities that aim to reduce mRNA levels. Unlike cytoplasmic siRNAs, which may be less effective when targeting the 5′UTR, splice-switching ASOs operate in the nucleus and modulate splicing with high specificity.^90–95^ Gapmer ASOs also act in the nucleus via RNase H-mediated degradation of pre-mRNA and mRNA,^91,92^ but lack isoform selectivity; because exon 2 is present in the pre-mRNA that gives rise to all the mRNA isoforms, gapmers would likely deplete both +E2 and -E2 transcripts, disrupting the dominant -E2 isoform in normal tissues. In contrast, splice-switching ASOs enable selective exon 2 skipping, knocking down the tumor-specific +E2 isoforms while sparing—actually promoting—the -E2 isoforms. This selectivity is expected to reduce on-target toxicity in normal tissues—which we confirmed using the HPDE immortalized pancreas cell line—and enhance therapeutic precision by exploiting the tumor-specific splicing landscape, in which exon 2 inclusion is driven by aberrantly high SRSF1 levels.

In summary, we described a novel mechanism of *AURKA* regulation, uncovering a splicing-dependent pathway to control oncogenic AURKA expression. We defined the SRSF1-AURKA-MYC circuit in PDAC, and demonstrated how splice-switching ASOs can effectively collapse this oncogenic circuit, inducing apoptosis in cancer cells. These results provide a strong foundation for ASO-based therapeutics that target oncogenic splicing programs, with broad implications for treating PDAC and other aggressive cancers.

## Limitations of the study

The experiments were carried out using *in vitro* model systems. Although organoids are powerful models, the next step will be to test the targeted ASO approach *in vivo*. Additionally, the ASO experiments involved transfection. We tested gymnotic delivery of *AURKA* ASO-A in PDAC organoids, but it was inefficient. Free uptake is highly dependent on both sequence and chemistry, as well as on the cells, so we plan to optimize these variables, and test other delivery approaches in the future. Lastly, ASO-A binds the AluJo-derived region of exon 2 and is predicted to hybridize with other genomic sites—mostly deeply intronic or intergenic Alu elements—so further optimization to minimize off-target effects may be necessary.

## RESOURCE AVAILABILITY

### Lead contact

Further information and requests for resources and reagents should be directed to and will be fulfilled by the lead contact, Adrian R. Krainer.

### Materials availability

Plasmids and unique/stable reagents generated in this study will be made available on request for scientific research.

### Data and code availability

- This study does not report original code.
- All data are available in the main text and the supplemental information or public datasets.
- Any additional information required to reanalyze the data reported in this paper is available from the lead contact upon request.

## ACKNOWLEDGMENTS

The Genotype-Tissue Expression (GTEx) Project was supported by the <u>Common Fund</u> of the Office of the Director of the National Institutes of Health, and by NCI, NHGRI, NHLBI, NIDA, NIMH, and NINDS. The data used for the analyses described in this manuscript were obtained from: the GTEx Portal on 03/18/2024. NCI Program Project Grant CA13106 Project 2 to A.R.K. Figures 1A, 7N, S3B-C were created using Biorender.com.

## AUTHOR CONTRIBUTIONS

Conceptualization, A.J.K. and A.R.K; methodology, A.J.K..; Investigation, A.J.K., L.J., G.Y.S., and L.W.; writing—original draft, A.J.K. and A.R.K.; writing—review & editing, A.J.K., L.J., L.W., and A.R.K.; funding acquisition, A.R.K.; resources, A.R.K.; supervision, A.R.K. CSHL Shared Resources used in this work were funded in part by NCI Cancer Center Support Grant 5P30CA045508.

## SUPPLEMENTAL INFORMATION

Document S1. Figures S1–S11 and Table S1-3

## METHOD DETAILS

### 2D cell lines and cell culture

HEK293T, SUIT2, and MiaPaca-2 cells were cultured in Dulbecco’s modified Eagle medium (DMEM) (Thermo Fisher, Waltham, MA) supplemented with 10% fetal bovine serum (Sigma-Aldrich, Burlington, MA) and 1% penicillin-streptomycin (Thermo Fisher). KLM1, BXPC3, and 786-O cells were cultured in RPMI- 1640 (Corning, Corning, NY) supplemented with 10% fetal bovine serum (Sigma-Aldrich) and 1% penicillin- streptomycin (Thermo Fisher). CAKI-1 cells were cultured with McCoy’s 5A (Modified) Medium (Thermo Fisher) supplemented with 10% fetal bovine serum (Sigma-Aldrich) and 1% penicillin-streptomycin (Thermo Fisher). HPDE cells were cultured in Keratinocyte SFM (1X) (Thermo Fisher) supplemented with Antibiotic- Antimycotic (100X) (Thermo Fisher). All cells were maintained at 37 °C

### Mouse organoid lines and culture

Mice KSC organoids were generated as previously described^6^.

### Human organoid lines and culture

Human PDAC organoids were obtained from the Organoids Shared Resource at CSHL and cultured at 37 °C as described^6,96,97^.

### RNA extraction, RT-PCR, and RT-qPCR

Total RNA was extracted with TRIzol (Thermo Fisher) according to the manufacturer’s protocol. RNA was treated with RQ1 DNase (Promega, Madison, WI) according to the manufacturer’s instructions, prior to cDNA synthesis. Oligo dT18-primed reverse transcription was carried out with ImProm-II reverse transcriptase (Promega). Semi-quantitative radioactive reverse transcription polymerase chain reaction (RT-PCR) was carried out in the presence of [α-^32^P] deoxycytidine triphosphate (Perkin Elmer, Waltham, MA) with primer sequences provided in Supplementary Table S2. PCR products were then separated by 5% native polyacrylamide gel electrophoresis (PAGE), detected with a Typhoon FLA7000 phosphorimager (GE, Boston, MA), and quantified using ImageJ software. For *AURKA*, all 5’UTR bands were captured and the sum of all +exon 2 isoforms was divided by the total intensity of all detected 5’UTR bands. For *SMN2*, PCR products were digested with DdeI restriction enzyme (NEB) for 1 h at 37 °C to separate *SMN1* and *SMN2* PCR products. Percent-spliced-in values for *AURKA* +E2 isoforms and *SMN2* were then calculated.

For RT-qPCR, Power Sybr Green Master Mix (Thermo Fisher) was used and cDNA was analyzed on a QuantStudio 6 Flex Real-Time PCR system (Thermo Fisher). Fold changes were calculated using the ΔΔCq method. Primer sequences are listed in Supplementary Table S2.

### Western blotting

Organoids and 2D cells were harvested and lysed in RIPA buffer (50 mM Tris, pH 8, 150 mM NaCl, 0.1% SDS, 1% NP-40, 0.5 % sodium deoxycholate) with cOmplete, mini protease inhibitor cocktail (MilliporeSigma, Burlington, MA) and Halt Phosphatase inhibitor cocktail (Thermo Fisher). Lysates were passed through a 25-gauge needle and cleared by centrifugation. Protein concentration was measured by Bradford assay (Bio-Rad, Hercules, CA) with BSA as a standard. A total of 30 μg of protein was separated via SDS-PAGE and transferred onto a nitrocellulose membrane. The following primary antibodies were used: AURKA (Cat# 14475), phosphor-T288 (Cat# 2914), phosphor-ERK1/2 (Cat# 4370), total-ERK1/2 (Cat# 4695), c-MYC (Cat# 5605) were purchased from Cell Signaling Technologies; Vinculin (Cat# sc- 73614) was purchased from Santa Cruz Biotechnologies; SRSF1 (CSHL clone AK96); antibody for mouse Aurka (Cat# 45-8900) purchased from Thermo Fisher. Antibodies were used with IRDye 800CW (Cat# 926-132210) or IRDye 680RD (Cat# 926-68071) secondary antibodies (LI-COR, Lincoln, NE) for western blotting, and the blots were imaged and quantified using an Odyssey Infrared Imaging System (LI-COR).

### Lentivirus production and transduction

Lentivirus was produced as previously described. Briefly, 7.5×10^6^ HEK293T cells were seeded in a 10-cm plate overnight. Target vector, psPAX2, and pMD2.G were co-transfected using Lipofectamine 3000 (Thermo Fisher) at 4:3:1 ratio, respectively. After 6 h, the medium was changed and the cells were incubated at 37 °C and 5% CO2. At 24 h and 48 h, cells were collected, pooled, filtered using a 0.45-μm syringe filter, and concentrated using a Lenti-X concentrator (Takara Bio, Japan). Cells or organoids were then transduced by spin inoculation at 900 × g for 45 min with 10 μg/ml polybrene (Sigma) supplemented to media. 24 h after infection, puromycin (Sigma) (5 μg/ml) was added to the cells for selection.

### Cloning procedures

Cloning of pCG-eGFP-AURKA-WT and pCG-eGFP-AURKA-D274N constructs was performed by inserting eGFP-AURKA cDNA sequence from pEGFP-Aurora A (Addgene #198402) into the pCG vector (Addgene #51476) using Gibson assembly. The D274N mutant was generated by site-directed mutagenesis using overlapping primers against pCG-eGFP-AURKA-WT.

All shRNA constructs were generated by the following protocol, using either pLKO.1-TRC (Addgene #10878) or Tet-pLKO-puro (Addgene #21915) as vectors. shRNA sequences for target genes were identified using (https://portals.broadinstitute.org/gpp/public/gene/search). Oligonucleotides were then boiled at 95 °C for 4 min in NEB 1× Buffer 2 (New England Biolabs, Ipswich, MA) and kept at room temperature for 2 h to anneal. Cloning vectors were linearized by restriction enzyme digestion using EcoR1- HF and AgeI-HF (NEB). Backbone and inserts were then ligated using T4 DNA ligase (NEB).

Cloning of *AURKA* cDNA 5’UTR reporters (Figure S5) was performed by inserting cDNA of each isoform into the pGL4.53 vector (Promega) using Gibson assembly.

Cloning of the *AURKA* 5’UTR minigene (Figure 4) and deletion mutants was performed as follows: *AURKA* 5’UTR (Gene ID: 6790- Chr:2 56,392,215- 56,388,202) was synthesized by GeneScript, spanning from the start of exon 1 through the first 5 nucleotides of exon 4, with flanking 5’ KpnI and 3’ NcoI restriction enzyme sites. The DNA fragment was inserted into the pGL4.53 firefly luciferase vector (Promega) via restriction sites, to generate pGL4-*AURKA*-5UTR-FL, into which all subsequent mutations were introduced. 3SS-Mut and ASO-res plasmids were generated by site-directed mutagenesis using overlapping primers along the mutation region. All deletion mutants were generated using the Q5 Site-Directed Mutagenesis Kit (NEB) with non-overlapping primers flanking each intron deletion.

### Dual-luciferase assay

All dual-luciferase assays were carried out using a Dual-luciferase Reporter Assay System (Promega) according to the manufacturer’s instructions. Briefly, 2 × 10^5^ SUIT2 cells were seeded on a 12-well plate. After 24 h, Firefly and pRL-SV40 Renilla Luciferase Control Reporter Vector (Promega) were co-transfected at 300 nM and 230 nM respectively, using Lipofectamine 3000 (Invitrogen). ASOs were also co-transfected where indicated at 30 nM concentration. After 24 h, luminescence for firefly and renilla was read using a SpectraMax i3 plate reader (Molecular Devices).

### Subcellular fraction of mRNA localization

SUIT2 cells were fractionated into cytoplasmic and nuclear fractions as described (https://rnajournal.cshlp.org/content/15/10/1896.long). Briefly, 4 × 10⁵ cells were seeded 6-well plates. After 24 h, cells were gently trypsinized and collected by centrifugation. The cell pellets were rinsed in 1× PBS, and the plasma membranes were lysed by resuspending the cells in 200 μl of ice-cold NP-40 lysis buffer (10 mM HEPES [pH 7.4], 0.5% NP-40, 10 mM NaCl, 3 mM MgCl₂) for 5 minutes. The lysate was then layered on top of 500 μl of a chilled sucrose cushion (24% sucrose in lysis buffer) and centrifuged at 2,300 rcf for 5 minutes at 4°C. The 200 μl supernatant (cytoplasmic fraction) was collected and mixed with 500 μl of TRIzol for RNA extraction. The nuclei pellet was gently rinsed with lysis buffer, resuspended in 100 μl of lysis buffer, and combined with 750 μl of TRIzol for RNA extraction.

### Chemical treatments

For KRAS^G12D^ inhibition RMC-9805 was purchased from MedChemExpress and dissolved in DMSO. SUIT2 cells were continually passaged in the presence of RMC-9805 at 10 μM for 1 month to generate the RMC- 980q5 resistant cell line. Bulk cell populations were used for downstream assays. RMC-9805 was used at a final concentration of 1 μM for functional assays.

For acute AURKA degradation, JB-170 was purchased from MedChemexpress and dissolved in DMSO. SUIT2 cells were treated with JB-170 at final concentration of 10 μM for the indicated times.

For AURKA kinase inhibition, VX-680 was purchased from MedChemExpress and dissolved in DMSO. SUIT2 cells were treated with VX-680 at final concentration of 1 μM for the indicated times.

For an apoptosis positive control, staurosporine (STS) was purchased from MedChemExpress and dissolved in DMSO. SUIT2 cells were treated with STS at final concentration of 10 μM for 16 h.

### Polysome profiling assay

Polysome profiling was adapted from^98^. Briefly, SUIT2 cells were grown to confluency in a 15-cm plate (∼ 2 x 10^7^ cells). Cells were treated with 50 μg/ml cycloheximide at 37 °C for 20 min. Cells were then washed with ice-cold PBS containing 50 μg/ml cycloheximide and lysed using 500 μl of polysome-extraction buffer (20 mM Tris pH 7.5, 5 mM MgCl2, 100 mM KCl, 0.5 % (v/v) nonyl phenoxylpolyethoxylethanol). Lysates were incubated on ice for 10 min and centrifuged at 13,000 × *g* for 10 min. 10-50% sucrose gradients were prepared using a BioComps Gradient Master by combining 10 % (w/v) and 50% sucrose solutions (20 mM Tris-HCL, pH 7.4, 100 mM KCl, 5 mM MgCl2) in a SW55 ultra-centrifuge tube. 300 μl of the cytoplasmic fraction was then layered on top of the gradient and samples were chilled at 4 °C for 1 h. Samples were then centrifuged at 4 °C in a Sorvall SW55 rotor at 36,000 r.p.m for 2 h. Fractions were then collected from the top using a BioComp piston gradient fractionator while measuring the OD at 254 nm. RNA was extracted from each fraction using Trizol and mRNA was analyzed using RT-qPCR.

### RNA stability assay

SUIT2 cells were seeded at 2 × 10^5^ cells in a 12-well plate. After 24 h, actinomycin D was added for each indicated time point at a final concentration of 5 μg/ml. The expression of each isoform was measured using specific primers by RT-qPCR. The half-life of each isoform was calculated using GraphPad Prism.

### Antisense oligonucleotide transfection

ASOs (uniformly modified with MOE sugars, phosphorothioate backbone, and 5-methyl cytosines) were purchased from Integrated DNA Technologies (IDT, Coralville, IA). 2D cells were transfected with the indicated amounts of ASO using Lipofectamine 3000. Cells were seeded and transfected at the same time. Organoid transfections were performed as described in.^97^

### RNA secondary structure prediction

RNA secondary structure of human *AURKA* 5’UTR (Chr:2 56,392,215- 56,388,202) was performed using RNAfold (v.2.6.3) through the ViennaRNA Web Server (http://rna.tbi.univie.ac.at/). The algorithm calculates the folding probability using a minimum free energy matrix with default parameters. In addition, the calculation included the partition function and base pairing probability matrix. RNA secondary structure prediction used the DNA sequence from *AURKA* 5’UTR and all of the predicted intron-deletion mutants. RNAcanvas (https://rna2drawer.app/) was used for structure annotations.

### Alu element identification and phylogenetic genome comparison

To identify Alu elements within *AURKA*, we used the UCSC Genome Browser (https://genome.ucsc.edu) and queried the RepeatMasker track, which annotates transposable elements across the genome. We filtered for Alu subfamilies in the human genome (GRCh38/hg38) to map their distribution within our regions of interest. To assess evolutionary conservation, we leveraged the UCSC Comparative Genomics tracks, using PhyloP and PhastCons scores to evaluate sequence conservation across species.

### Apoptosis assay

To measure apoptosis, Guava Nexin Reagent (Cytek) was used according to the manufacturer’s instructions. Briefly, cell-culture supernatant was transferred to 15-ml conical tubes, before the cells were washed with 1 × PBS, to capture floating cells. Cells were then trypsinized and neutralized with original supernatant for each condition. Cells were then transferred to a 15-ml conical tube and pelleted by centrifugation at 300 × *g* for 5 min. Cell pellets were then resuspended with culture media to a concentration of 10^6^ cells/ml. 100 μl of each sample was pipetted into a 96-well plate and then 100 μL of Guava Nexin Reagent was added to each well. Samples were incubated for 20 min at room temperature in the dark and were then acquired on a Guava EasyCyte HT-BG benchtop flow cytometer (Cytek).

### Cell viability assay

To measure cell viability, CellTiter-Glo 3D was used according to the manufacturer’s instructions. Briefly, 5000 organoids or cells were seeded in opaque-walled 96-well plates and luminescence was read at 72 h using CellTiter-Glo 3D (Promega). Luminescence reading of organoids after 72 h was normalized to reading immediately after seeding.

### Whole-mount organoid staining and immunofluorescence

Organoid immunofluorescence staining was performed as previously described, with minor adjustments^97^. Briefly, Organoids were transfected with ASO as described above. After 72 h, organoids were harvested by incubation with Cell Recovery solution for 1 h on a horizontal rocker at 4 °C. After fixation, organoids were incubated with primary antibody MOEC (Rockland, 1:1000) overnight. Organoids were then incubated with Alexa Fluor Plus 488 (Thermo Fisher) diluted in blocking buffer with DAPI and Phallodin-Atto 565 (Sigma). Organoids were then mounted on concave microscope slides, and imaged on a Zeiss LSM 780 confocal microscope.

### RNA splicing analysis

ΔPSI and FDR values of splicing changes were obtained using PSI-Sigma v1.9j with parameters: “–gtf Mus_musculus.GRCm38.100.stringtie.sorted.gtf –nread 20 –type 1 –fmode 3 –irrange 5”. StringTie v2.1.4 was used to generate the “Mus_musculus.GRCm38.100.stringtie.sorted.gtf” file for building the splicing database of PSI-Sigma.

### Analysis of public datasets

The association of *AURKA* mRNA levels in pancreas, kidney, and liver samples with clinical prognosis was determined using UCSC Xena browser (https://xenabrowser.net/) based on TCGA data. The comparisons of *AURKA* mRNA levels between pancreatic tumor and normal, kidney tumor and normal, and liver tumor and normal were also performed using UCSC Xena browser based on TCGA and GTEx data. The correlation between *SRSF1* and *AURKA* mRNA levels in pancreas tumors and cell lines was determined using UCSC Xena browser based on TCGA and DepMap Portal (https://depmap.org/portal), respectively.

### Identification of splicing cis-regulatory sequences

To identify potential SRSF1 ESE sequences, we used *AURKA* exon 2 DNA sequence as input for ESEfinder3.0 (https://esefinder.ahc.umn.edu/cgi-bin/tools/ESE3/esefinder.cgi). SF2/ASF (a.k.a. SRSF1) and SF2/ASF (IgM-BRCA1) matrices were used to identify predicted ESEs. To determine 5’ and 3’ splice site strengths, we used MaxEntScan::score5ss (http://hollywood.mit.edu/burgelab/maxent/Xmaxentscan_scoreseq.html) and MaxEntScan::score3ss (http://hollywood.mit.edu/burgelab/maxent/Xmaxentscan_scoreseq_acc.html).

## QUANTIFICATION AND STATISTICAL ANALYSIS

Data for quantification analyses are represented as mean ± SD. The significance of differences was examined by Student’s t test (*P ≤ 0.05, **P ≤ 0.01, ***P ≤ 0.001, ****P ≤ 0.0001) or one-way ANOVA followed by Dunnet’s multiple-comparison test. The statistical analyses for all experiments were performed with GraphPad Prism 9 software (https://www.graphpad.com/scientific-software/prism/). The exact value of n, what n represents, and the statistical details of each experiment are described in the figure legends or tables.

## Authors Contributions

**YS**- designing the experiment, writing software, testing participants, analyzing data, writing the paper

**SG**- testing participants, analyzing data, writing the paper

**GK-** designing the experiment, managing resources, analyzing data, writing the paper

---

Figure S1. Mouse *Aurka* and human *AURKA* 5’UTRs undergo alternative splicing and comprise conserved SRSF1 ESE motifs.

(A and B) Schematic of mouse *Aurka* (A) and human *AURKA* (B) genes, highlighting the 5’UTRs.

(C) Sequence alignment of SRSF1 ESE motifs predicted by ESEfinder 3.0 found in exon 2 across the indicated species.

(D) GTEx transcript browser view (https://www.gtexportal.org/home/transcriptPage) of *AURKA* isoform expression across normal tissues. Pancreas is indicated by red box.

Figure S2. AluSp and AluSq2 sequences are nearly perfect reverse complements

(A) Sequence alignment of AluSp and AluSq2 elements. The positions of exon 2 and exon 3 are indicated.

Figure S3. AluSp and AluSq2 are inverted Alu elements, potentially form a dsRNA structure in *AURKA* pre-mRNA, and result in exonization of exons 2 and 3 in the *AURKA* 5’UTR

(A) Secondary structure of the full-length *AURKA* 5’UTR predicted by RNAfold. The locations of exons, introns, AluSp, and AluSq2 are indicated.

(B) Schematic of the size, position, and orientation of AluJo, AluSp, and AluSq2 elements in relation to exon 2 and exon 3 (top). Potential secondary structure of AluSp and AluSq2 (bottom).

(C) Schematic and sequence alignments for exon 2 (top) and exon 3 (bottom) 3’ and 5’ splice sites. The sequences are aligned to the ancestral Alu sequences from the respective subfamilies. The positions of 3’ and 5’ splice sites are indicated in red, with the corresponding splice-site strength scores from MaxEntScore::3SS and MaxEntScore::5SS.

Figure S4. *AURKA* mRNA is upregulated, associated with poor survival outcomes, and highly correlated with *SRSF1* mRNA levels in PDAC.

(A) Violin plot of *AURKA* mRNA expression in normal and tumor samples from TCGA and GTEx (n = 167 for normal and n = 183 for tumor). Data from UCSC Xena.

(B) Kaplan-Meier survival analysis of patients with pancreatic adenocarcinoma from TCGA. Group cut-offs were set at the *AURKA* median fpkm expression levels (n = 92 for low expression and n = 91 for high expression). Data from UCSC Xena.

(C) Correlation of mRNA expression levels of *SRSF1* vs. *AURKA* of pancreatic adenocarcinoma patients from TCGA (n = 179) using simple linear regression. Data from UCSC Xena.

(D) Correlation of mRNA expression levels of *SRSF1* vs. *AURKA* of pancreas-cancer-derived cell lines (n

= 54) using simple linear regression. Data from DepMap portal https://depmap.org/portal.

(E) RT-qPCR correlation of total *SRSF1* mRNA vs. total *AURKA* mRNA normalized to *RPS13* mRNA from patient-derived organoids, using simple linear regression.

Data shown as mean ± SD. ****p ≤ 0.0001 Student’s t test (A), log rank (Mantel-Cox) test (B), and correlation and simple linear regression (C, D, and E).

Figure S5. Exon-2-inclusion isoforms associate with heavier polysomes.

(A) Schematic of *AURKA* 5’UTR cDNA reporters. Each reporter has the corresponding RefSeq accession number listed below. For simplicity, we arbitrarily named each isoform “ISO1-5”.

(B) RLU quantification of dual-luciferase assay from HEK293T cells after Lipofectamine 3000 transfection of *AURKA* 5’UTR cDNA reporters from (B). Each reporter was transfected at a concentration of 500 ng. Firefly luciferase activity was normalized to Renilla luciferase activity. Each condition was normalized to EV (n = 3 biological replicates). RLU, relative luciferase units; EV, empty vector.

(C) Schematic of Actinomycin D RNA-stability assay.

(D) RNA-stability assay of endogenous *AURKA* isoforms in SUIT2 cells harvested at the indicated times after actinomycin D treatment (C). mRNAs were quantified using RT-qPCR, normalized to *RPS13*, and expressed as a percentage of the levels at time 0 h (n = 3 biological replicates).

(E) Polysome profile of endogenous *AURKA* isoforms in SUIT2 cells. mRNA from each fraction was quantified using RT-qPCR, normalized to *RPS13*, and expressed as a percentage of the total mRNA from the sum of all fractions for each isoform (n = 3 biological replicates). LMW, low molecular weight; HMW, high molecular weight.

(F-H) Area under the curve (AUC) quantification of (E) for polysome fractions 9-15 (F), 9-12 (G), and 12-15

(H) (n = 3 biological replicates).

Data shown as as mean ± SD. ns = non-significant, **p ≤ 0.01, ***p ≤ 0.001 one-way ANOVA (B) or Student’s t test (F, G, H).

Figure S6. Splicing and localization analysis of the *AURKA* 5’UTR minigenes.

(A) Snapgene snapshot of FL *AURKA* 5’UTR minigene, highlighting the positions of the primers, exons, and Alu elements.

(B) Radioactive RT-PCR of *AURKA* 5’UTR minigene mRNA from SUIT2 cells transfected with Lipofectamine 3000 and the indicated vectors (top). Radioactive RT-PCR of *Renilla* mRNA (bottom).

(C) Radioactive RT-PCR of *AURKA* 5’UTR minigene, *MTRNR2*, and *NEAT1* RNA of SUIT2 cells after Lipofectamine 3000 transfection of minigene vectors and subsequent subcellular fractionation. W, whole- cell fraction; C, cytoplasmic fraction; N, nuclear fraction.

Figure S7. Predicted RNA secondary structure of *AURKA* 5’UTR minigenes.

(A-G) Secondary structure of *AURKA* 5’UTR intron deletion minigene mutants, as predicted by RNAfold. Structures were analyzed using RNAcanvas. The positions of exon 2, exon 3, AluSp, AluSq2, and the length of the dsRNA region are indicated for each intron-deletion mutant.

Figure S8. High concentration of VX-680 and extended treatment time leads to reductions of AURKA, SRSF1, and MYC proteins.

(A) Western blotting of AURKA, SRSF1, MYC, and VINC after VX-680 [10 μM] treatment of SUIT2 cells for 72 h.

(B) Relative protein expression quantification compared to SUIT2 DMSO-treated cells for AURKA (left), SRSF1 (middle), and MYC (right). Band intensity was normalized to the VINC loading control. The VX-680 samples were then normalized to the DMSO control samples (n = 3 biological replicates).

Data are represented as mean ± SD. *p ≤ 0.05, **p ≤0.01 Student’s t test (B).

Figure S9. Identification of ASO-A.

(A) Schematic of *AURKA* exon 2 3’ss-targeting ASOs.

(B) Western blotting of AURKA, SRSF1, MYC, and VINC after Lipofectamine 3000 ASO transfection at 100 nM for 48 h. 3, ASO-3; 3.1, ASO-3.1; 3.2, ASO-3.2; 3.3, ASO-3.3.

(C) Radioactive RT-PCR of *AURKA* 5’ UTR from SUIT2 cells after Lipofectamine 3000 transfection with ASOs as in (B).

(D) PSI quantification of +E2 isoforms from (C) (n = 3 biological replicates).

(E) RT-qPCR of total *AURKA* mRNA from (C). mRNA expression normalized to *RPS13* (n = 3 biological replicates).

(F) CellTiter-Glo cell viability assay of SUIT2 cells transfected with ASO as in (C). Each condition was normalized to the CellTiter-Glo reading immediately after cell seeding (Day 0) (n = 3 biological replicates).

(G) Radioactive RT-PCR of *AURKA* 5’UTR from SUIT2 cells after Lipofectamine 3000 transfection of ASO-A at 65 nM for the indicated times

(H) PSI quantification of +E2 isoforms from (G) (n = 3 biological replicates).

(I) RT-qPCR of total *AURKA* mRNA from (G). mRNA normalized to *RPS13* (n = 3 biological replicates).

(J) Western blotting of AURKA, SRSF1, MYC, and VINC after ASO transfection as in (G). Relative fold change for each protein compared to NTC is provided. Bands were normalized to VINC and then normalized to NTC. FC, fold change.

(K) Radioactive RT-PCR of *AURKA* 5’UTR from SUIT2 cells after Lipofectamine 3000 of N, S (65 nM), ASO-6 (65 nM), or ASO-A (30 nM). 6, ASO-6; A, ASO-A.

(L) PSI quantification of +E2 isoforms from (K) (n = 3 biological replicates).

(M) RT-qPCR of total *AURKA* mRNA from (K). mRNA normalized to *RPS13* (n = 3 biological replicates).

(N) Western blotting of AURKA, SRSF1, MYC, and VINC from SUIT2 cells after transfection of ASO as in

(I). Relative fold change for each protein compared to NTC is provided. Bands were normalized to VINC and then normalized to NTC.

Data are represented as mean ± SD. ns = non-significant, *p ≤ 0.05, ***p ≤ 0.001, ****p ≤ 0.0001 one-way ANNOVA (D, E, F, H, I, L, and M).

Figure S10. ASO-A inhibits human PDAC organoid growth and collapses the SRSF1-AURKA-MYC axis

(A) Radioactive RT-PCR of *AURKA* 5’UTR from hT1 organoids after Lipofectamine 3000 transfection at 125 nM for 72 h. NTC, non-treated control (Lipofectamine 3000 only); SC, scrambled control; A, ASO-A.

(B-E) PSI quantification of +E2 isoforms (B) and RT-qPCR of total *AURKA* (C), *SRSF1* (D), and *MYC* (E) mRNA from (A). mRNA expression normalized to *RPS13* (C-E) (n = 3 biological replicates).

(F) Western blotting of AURKA, SRSF1, MYC, and VINC after ASO treatment of hT1 organoids as in (A).

(G) CellTiter-Glo cell-viability assay of hT1 organoids transfected with ASO as in (A). Each condition was normalized to the CellTiter-Glo reading immediately after cell seeding (Day 0) and then normalized to NTC.

(H) Radioactive RT-PCR of *SMN2* (top), and *RPS13* (bottom) from HPDE (left) and SUIT2 (right) cells after Lipofectamine 3000 transfection at 30 nM for 48 h. PSI for *SMN2* exon 7 is provided. Nus. Nusinersen.

Data are represented as mean ± SD. ns = non-significant; *p ≤ 0.05; ***p ≤ 0.001, ****p ≤ 0.0001 one-way ANOVA (C, D, E, G).

Figure S11. ASO-A induces exon 2 skipping in multiple cancer cell lines.

(A) Violin plot (left) of *AURKA* mRNA expression in normal and tumor kidney samples from TCGA and GTEx (n = 28 for normal and n = 884 for tumor). Kaplan-Meier (right) survival analysis of patients with renal cell carcinoma from TCGA. Group cutoffs were set at the *AURKA* median fpkm expression level (n = 437 for low expression and n = 445 for high expression). Data from UCSC Xena.

(B) Violin plot (left) of *AURKA* mRNA expression in normal and tumor liver samples from TCGA and GTEx (n = 110 for normal and n = 431 for tumor). Kaplan-Meier (right) survival analysis of patients with hepatocellular carcinoma from TCGA. Group cutoffs were set at the *AURKA* median fpkm expression level (n = 184 for low expression and n = 184 for high expression). Data from UCSC Xena.

(C) Radioactive RT-PCR of *AURKA* 5’UTR in PDAC (left), RCC (middle), and HCC (right) cell lines after Lipofectamine 3000 transfection of ASO at 65 nM for 48 h. SC, scrambled control; A, ASO-A; RCC, Renal Cell Carcinoma; HCC, Hepatocellular carcinoma.

(D) PSI quantification of +E2 isoforms from (C). Each condition is compared to SC (n = 3 biological replicates).

(E) Western blotting of AURKA, SRSF1, MYC, and VINC after ASO transfection as in (C). A, AURKA; S, SRSF1; M, MYC; V, VINC.

(F) CellTiter-Glo cell-viability assay after ASO transfection as in (C) (n = 3 biological replicates).

Data are represented as mean ± SD. ns = non-significant, *p ≤ 0.05, **p ≤ 0.01, ***p ≤ 0.001, ****p ≤ 0.0001 Student’s t test (A, B, D, F), Log rank (Mantel-Cox) test (A and B).

