## Supplemental Document S1 for "Splice-switching ASOs targeting an Alu-derived exon in the *AURKA* 5’UTR collapse an SRSF1-AURKA-MYC oncogenic circuit in pancreatic cancer"

Figure S1:

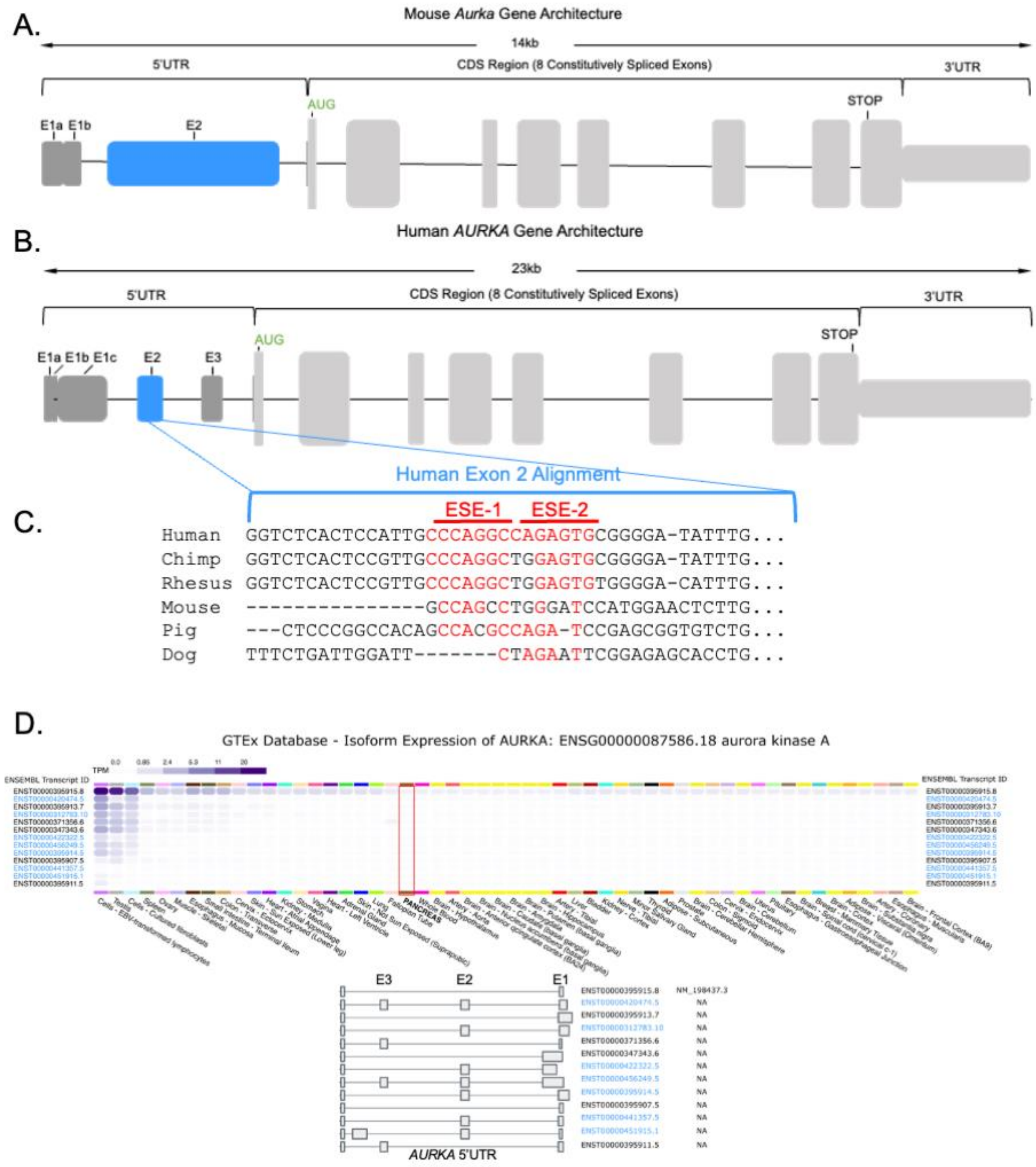

A.
Mouse *Aurka* Gene Architecture
14kb
5'UTR
CDS Region (8 Constitutively Spliced Exons)
3'UTR
AUG
STOP
E1a E1b
E2
B.
Human *AURKA* Gene Architecture
23kb
5'UTR
CDS Region (8 Constitutively Spliced Exons)
3'UTR
AUG
STOP
E1a E1b E1c
E2
E3
Human Exon 2 Alignment
C.
ESE-1
ESE-2
Human GGTCTCACTCCATTGCCCAGGCCAGAGTGCGGGGA-TATTTG...
Chimp GGTCTCACTCCGTTGCCCAGGCTGGAGTGCGGGGA-TATTTG...
Rhesus GGTCTCACTCCGTTGCCCAGGCTGGAGTGTGGGGA-CATTTG...
Mouse ---------------GCCAGCCTGGGATCCATGGAACTCTTG...
Pig ---CTCCCGGCCACAGCCACGCCAGA-TCCGAGCGGTGTCTG...
Dog TTTCTGATTGGATT-------CTAGAATTCGGAGAGCACCTG...
D.
GTEx Database - Isoform Expression of AURKA: ENSG00000087586.18 aurora kinase A
TPM
ENSEMBL Transcript ID
PANCREAS
E3
E2
E1
*AURKA* 5'UTR

Figure S2:

A.

```
                              5' - Exon 2 - 3'
AluSp (-)    1    GGCCGGGCGCGGTGGCTCATGCCCGTAATCCCAGCATTTTCGGAGGCCGAGGTGGGTGGA   60
                  ||||||||  ||||||| |  || |||||||||||||||| ||||||||||||||| |||
AluSq2(+)    1    GGCCGGGCATGGTGGCTTACACCTGTAATCCCAGCATTTTGGGAGGCCGAGGTGGGCGGA   60

AluSp (-)   61    TCACCTGAGGTCGGGAGTTCCAGACCAGCCCTACCAACATAGAGAAACCCTGTCTCTACT   120
                  ||||||||||||  |||||| |||||||||   ||||||| | ||||||| |||||||||
AluSq2(+)   61    TCACCTGAGGTCAAGAGTTCGAGACCAGCCTGGCCAACATGGTGAAACCCCGTCTCTACT   120

AluSp(-)   121    AAAAATACAAAA-TCAGCTGGGCGTGGTGGTGCATGT---CTGTAATCCCAGCTACTCTG   176
                  |||||||||||| | ||||| ||||||||  |||  ||   |||||||||||||||||| |
AluSq2(+)  121    AAAAATACAAAACTTAGCTGAGCGTGGTG--GCA-GTCACCTGTAATCCCAGCTACTCGG   177

AluSp (-)  177    GAGGCTGAGGCAGGAGAATTGCTTGAACCCAGGAGGTGGAGGTTGCGGGTGAGCCGAGAT   236
                  |||||||||||||||||||   |||  ||||||||| |  ||||||||  |||||| |||||
AluSq2(+)  178    GAGGCTGAGGCAGGAGAATCACTTAAACCCAGGAAGCAGAGGTTGC-AGTGAGCTGAGAT   236

AluSp (-)  237    TGCGCCATTGCACTCCAGCCTGGGCAACAAGAGC-AAACTCTATCTCAAAAAAAAAAAAA   295
                    | |||||||||||||||||  |  | |||||| | |||   |||||||| || |||| 
                     3' - Exon 3 - 5'
AluSq2(+)  237    CACACCATTGCACTCCAGCCTATGGGAAAAGAGCGAGACTTCGTCTCAAAACAATAAAAT   296

AluSp (-)  296    AA   297
                  ||
AluSq2(+)  297    AA   298
```

Figure S3:

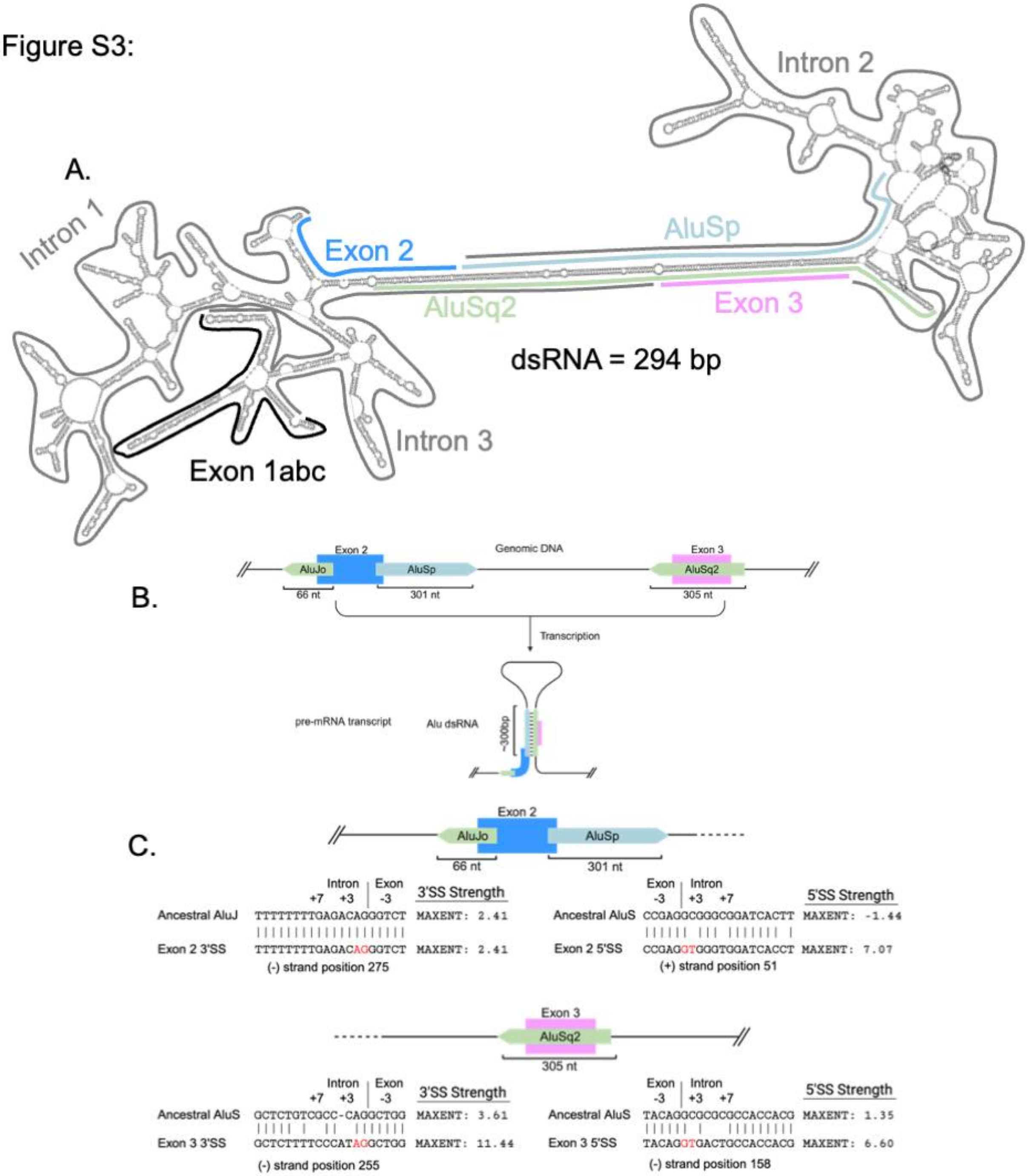
A.
Intron 1
Intron 2
Intron 3
Exon 1abc
Exon 2
AluSp
AluSq2
Exon 3
dsRNA = 294 bp
B.
Genomic DNA
Exon 2
AluJo
AluSp
66 nt
301 nt
Exon 3
AluSq2
305 nt
Transcription
pre-mRNA transcript
Alu dsRNA
~300bp
C.
Exon 2
AluJo
AluSp
66 nt
301 nt
Intron | Exon
+7 +3 -3
3'SS Strength
Ancestral AluJ TTTTTTTTGAGACAGGGTCT MAXENT: 2.41
Exon 2 3'SS TTTTTTTTGAGACAGGGTCT MAXENT: 2.41
(-) strand position 275
Exon | Intron
-3 +3 +7
5'SS Strength
Ancestral AluS CCGAGGCGGGCGGATCACTT MAXENT: -1.44
Exon 2 5'SS CCGAGGTGGGTGGATCACCT MAXENT: 7.07
(+) strand position 51
Exon 3
AluSq2
305 nt
Intron | Exon
+7 +3 -3
3'SS Strength
Ancestral AluS GCTCTGTCGCC-CAGGCTGG MAXENT: 3.61
Exon 3 3'SS GCTCTTTTCCCATAGGCTGG MAXENT: 11.44
(-) strand position 255
Exon | Intron
-3 +3 +7
5'SS Strength
Ancestral AluS TACAGGCGCGCCACCACG MAXENT: 1.35
Exon 3 5'SS TACAGGTGACTGCCACCACG MAXENT: 6.60
(-) strand position 158

Figure S4:

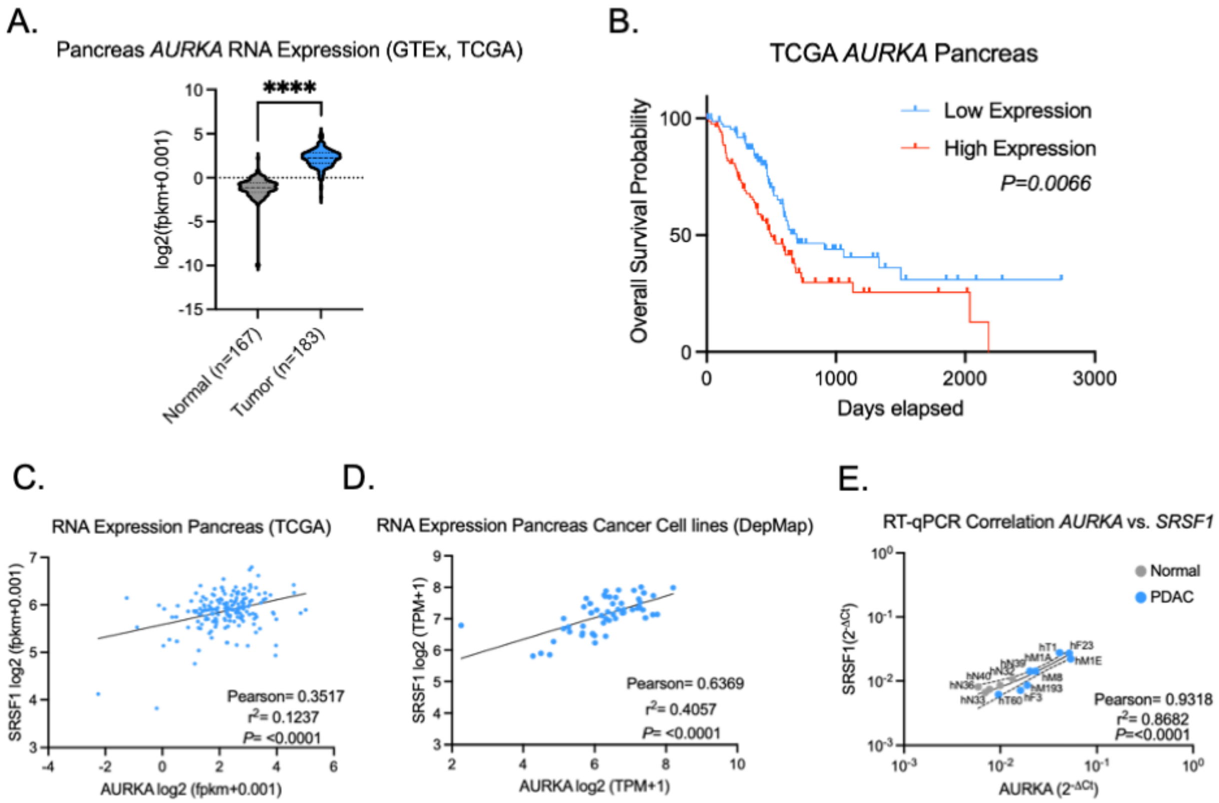
A.
Pancreas AURKA RNA Expression (GTEx, TCGA)
****
log2(fpkm+0.001)
Normal (n=167)
Tumor (n=183)
B.
TCGA AURKA Pancreas
Low Expression
High Expression
P=0.0066
Overall Survival Probability
Days elapsed
C.
RNA Expression Pancreas (TCGA)
SRSF1 log2 (fpkm+0.001)
Pearson= 0.3517
r2= 0.1237
P= <0.0001
AURKA log2 (fpkm+0.001)
D.
RNA Expression Pancreas Cancer Cell lines (DepMap)
SRSF1 log2 (TPM+1)
Pearson= 0.6369
r2= 0.4057
P= <0.0001
AURKA log2 (TPM+1)
E.
RT-qPCR Correlation AURKA vs. SRSF1
Normal
PDAC
SRSF1 (2-ΔCt)
Pearson= 0.9318
r2= 0.8682
P=<0.0001
AURKA (2-ΔCt)

Figure S5:

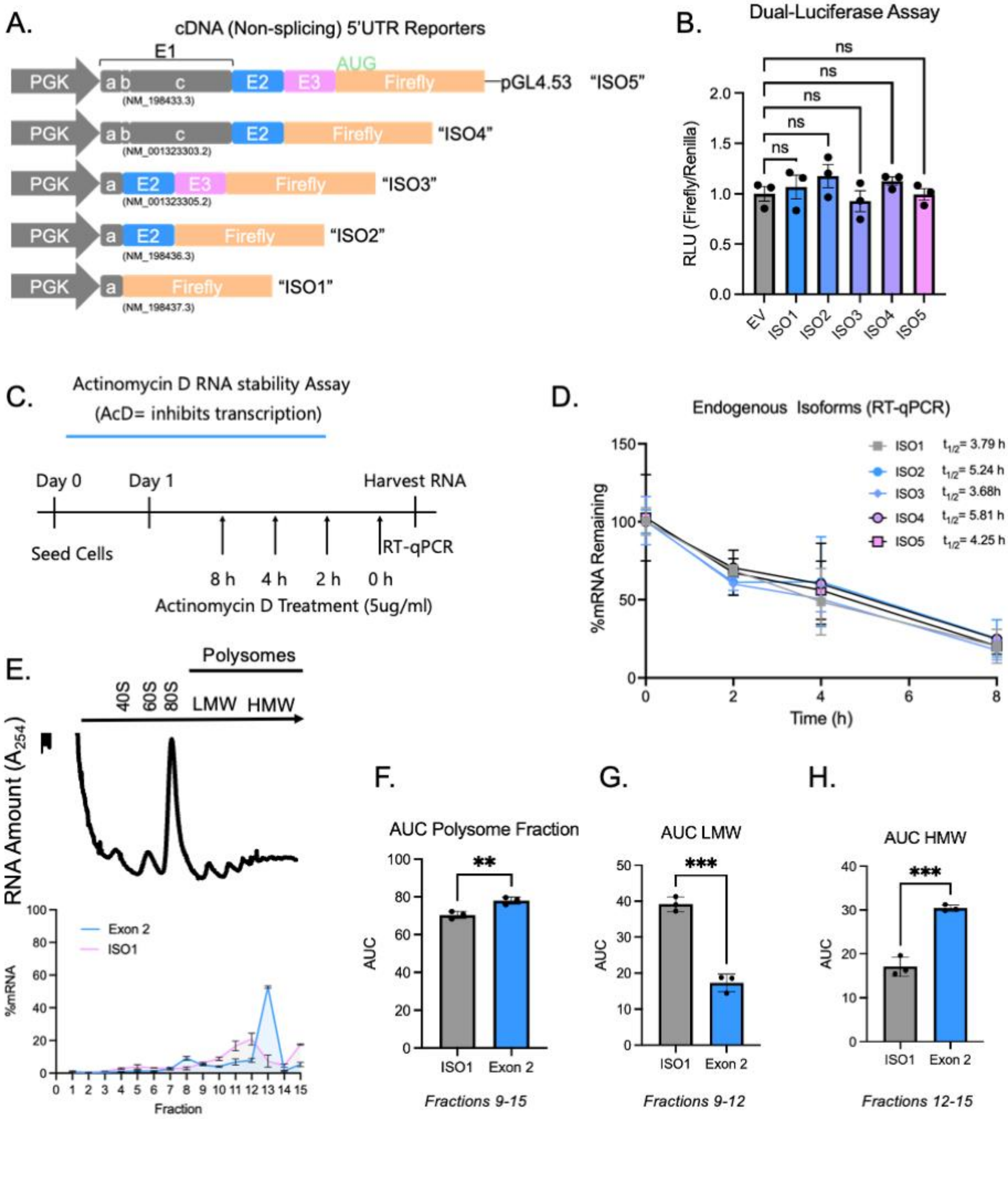

A.
cDNA (Non-splicing) 5'UTR Reporters
PGK
E1
AUG
Firefly
pGL4.53
"ISO5"
"ISO4"
"ISO3"
"ISO2"
"ISO1"
(NM_198433.3)
(NM_001323303.2)
(NM_001323305.2)
(NM_198436.3)
(NM_198437.3)
B.
Dual-Luciferase Assay
RLU (Firefly/Renilla)
ns
EV
ISO1
ISO2
ISO3
ISO4
ISO5
C.
Actinomycin D RNA stability Assay
(AcD= inhibits transcription)
Day 0
Day 1
Harvest RNA
Seed Cells
RT-qPCR
8 h
4 h
2 h
0 h
Actinomycin D Treatment (5ug/ml)
D.
Endogenous Isoforms (RT-qPCR)
%mRNA Remaining
Time (h)
ISO1 t1/2= 3.79 h
ISO2 t1/2= 5.24 h
ISO3 t1/2= 3.68h
ISO4 t1/2= 5.81 h
ISO5 t1/2= 4.25 h
E.
Polysomes
40S
60S
80S
LMW
HMW
RNA Amount (A254)
Exon 2
ISO1
%mRNA
Fraction
F.
AUC Polysome Fraction
AUC
Exon 2
Fractions 9-15
G.
AUC LMW
Fractions 9-12
H.
AUC HMW
Fractions 12-15

Figure S6:

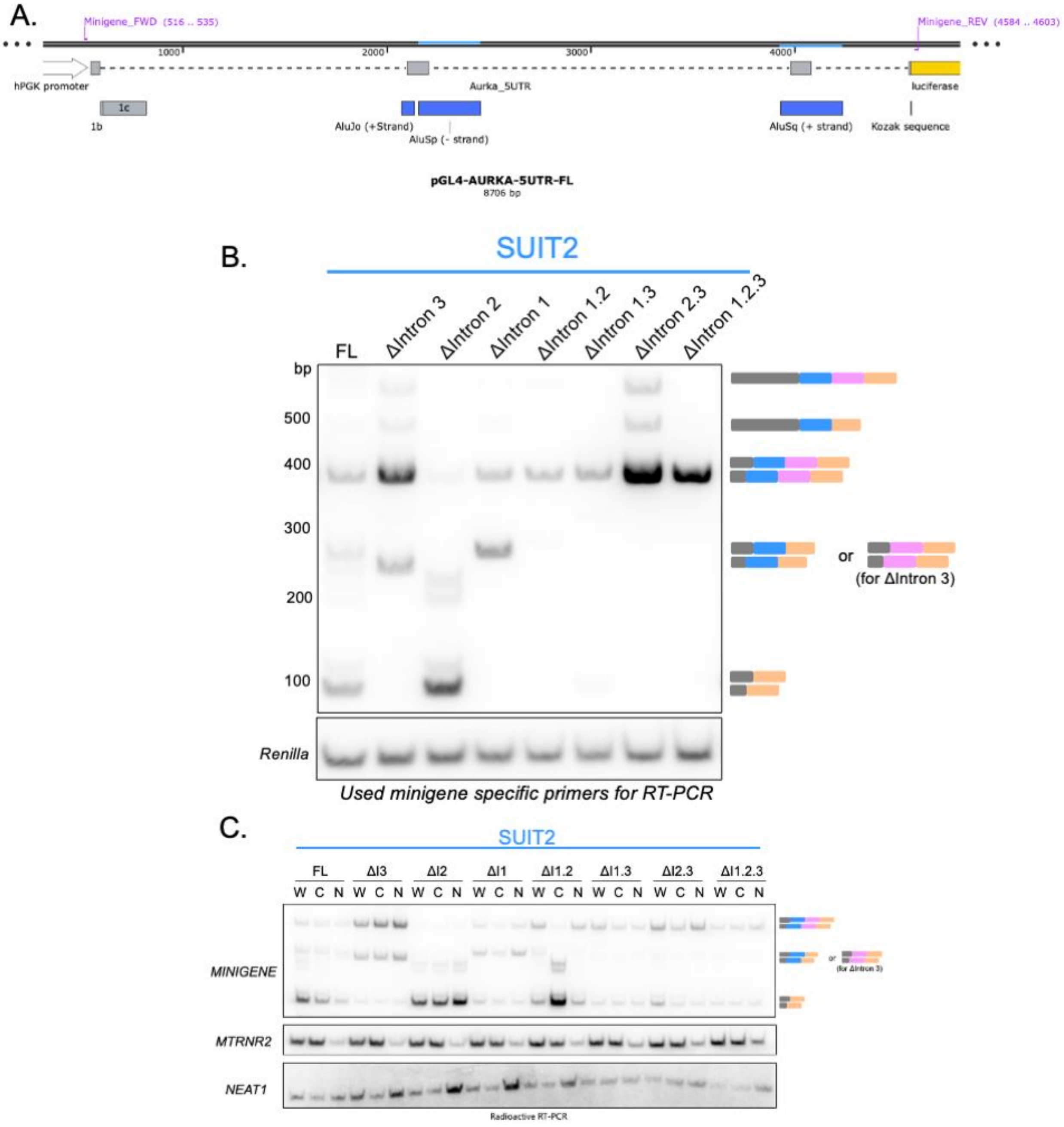
A.
Minigene_FWD (516 .. 535)
Minigene_REV (4584 .. 4603)
1000
2000
3000
4000
hPGK promoter
Aurka_5UTR
luciferase
1c
1b
AluJo (+Strand)
AluSp (- strand)
AluSq (+ strand)
Kozak sequence
pGL4-AURKA-5UTR-FL
8706 bp
B.
SUIT2
FL
ΔIntron 3
ΔIntron 2
ΔIntron 1
ΔIntron 1.2
ΔIntron 1.3
ΔIntron 2.3
ΔIntron 1.2.3
bp
500
400
300
200
100
or
(for ΔIntron 3)
Renilla
Used minigene specific primers for RT-PCR
C.
SUIT2
FL
ΔI3
ΔI2
ΔI1
ΔI1.2
ΔI1.3
ΔI2.3
ΔI1.2.3
W C N W C N W C N W C N W C N W C N W C N W C N
MINIGENE
MTRNR2
NEAT1
Radioactive RT-PCR

Figure S7:

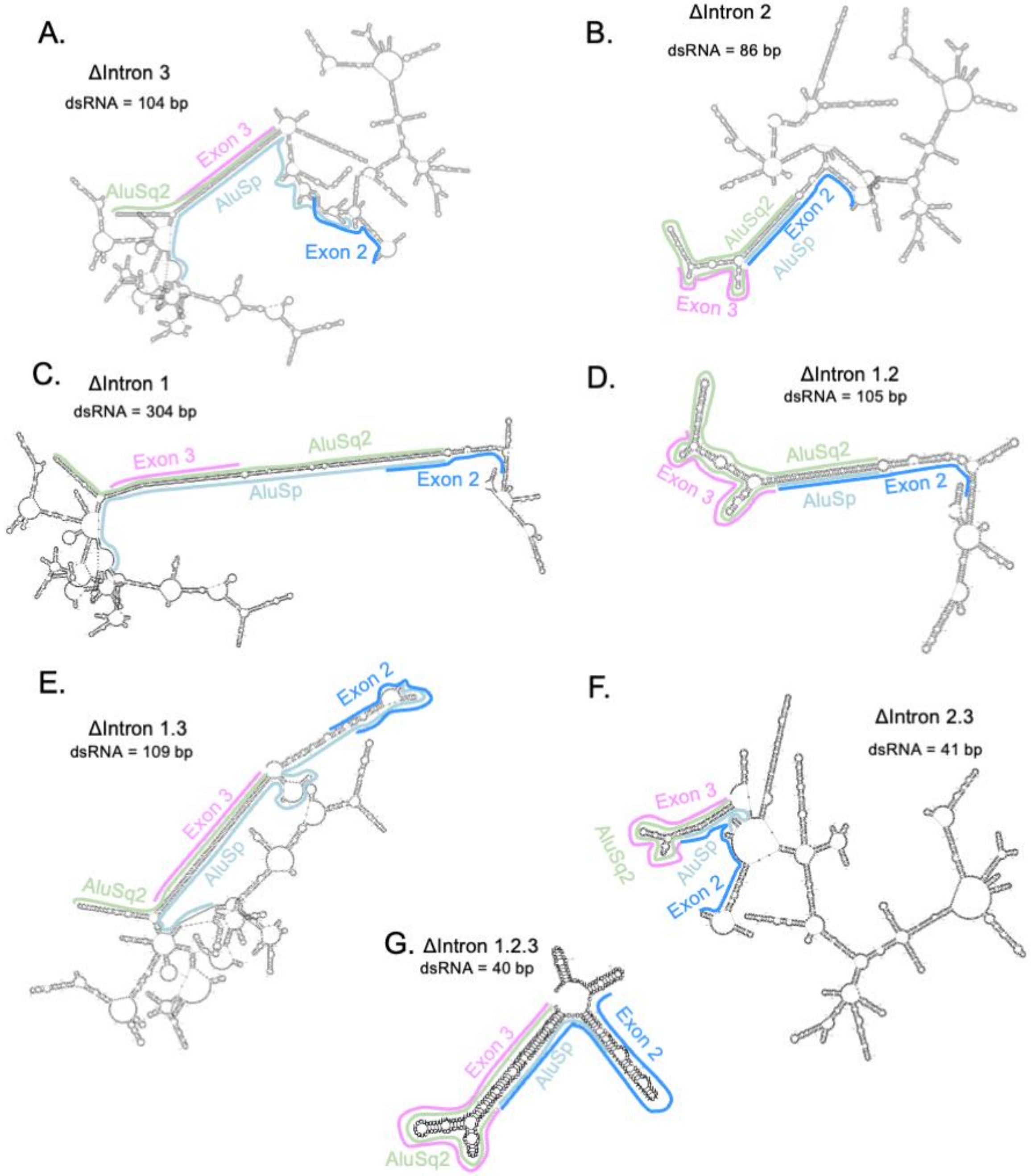
A.
ΔIntron 3
dsRNA = 104 bp
Exon 3
AluSq2
AluSp
Exon 2
B.
ΔIntron 2
dsRNA = 86 bp
AluSq2
Exon 2
AluSp
Exon 3
C.
ΔIntron 1
dsRNA = 304 bp
Exon 3
AluSq2
AluSp
Exon 2
D.
ΔIntron 1.2
dsRNA = 105 bp
AluSq2
Exon 3
AluSp
Exon 2
E.
ΔIntron 1.3
dsRNA = 109 bp
Exon 2
Exon 3
AluSp
AluSq2
F.
ΔIntron 2.3
dsRNA = 41 bp
Exon 3
AluSq2
AluSp
Exon 2
G. ΔIntron 1.2.3
dsRNA = 40 bp
Exon 3
AluSp
Exon 2
AluSq2

Figure S8:

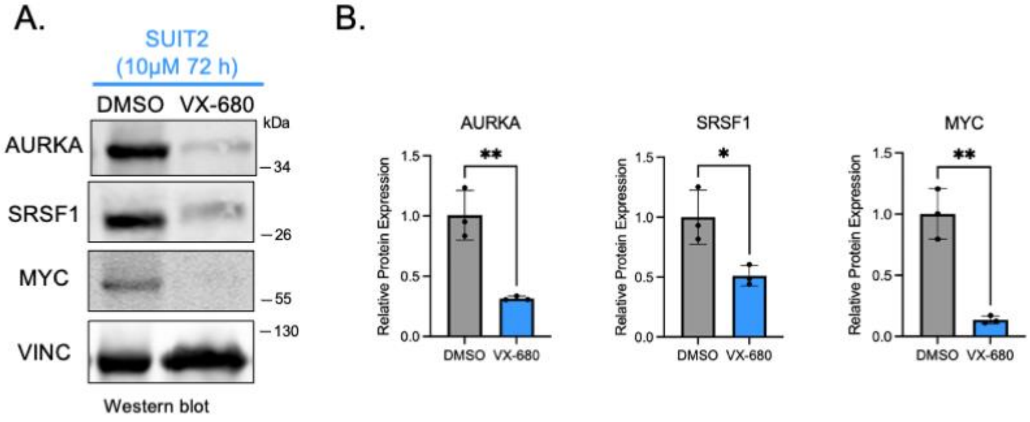
A.
SUIT2
(10μM 72 h)
DMSO
VX-680
kDa
AURKA
34
SRSF1
26
MYC
55
130
VINC
Western blot
B.
AURKA
SRSF1
MYC
Relative Protein Expression
1.5
1.0
0.5
0.0
**
*
**
DMSO
VX-680

Figure S9:

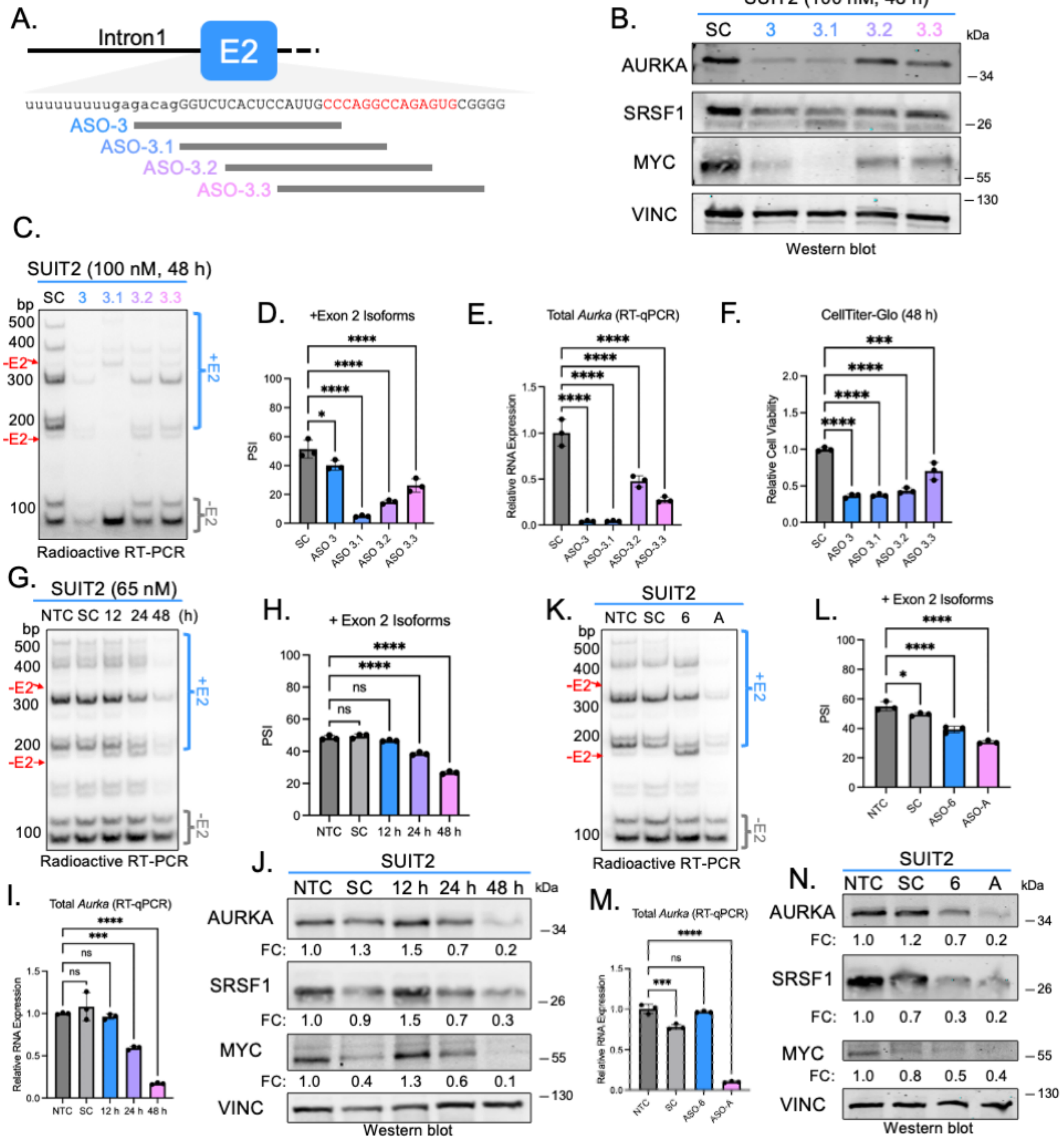
A.
Intron1
E2
uuuuuuuuugagacagGGUCUCACUCCAUUGCCCAGGCCAGAGUGCGGGG
ASO-3
ASO-3.1
ASO-3.2
ASO-3.3
B.
SUIT2 (100 nM, 48 h)
SC 3 3.1 3.2 3.3
kDa
AURKA
SRSF1
MYC
VINC
34
26
55
130
Western blot
C.
SUIT2 (100 nM, 48 h)
SC 3 3.1 3.2 3.3
bp
+E2
-E2
Radioactive RT-PCR
D.
+Exon 2 Isoforms
PSI
E.
Total Aurka (RT-qPCR)
Relative RNA Expression
F.
CellTiter-Glo (48 h)
Relative Cell Viability
G.
SUIT2 (65 nM)
NTC SC 12 24 48 (h)
Radioactive RT-PCR
H.
+ Exon 2 Isoforms
PSI
NTC SC 12 h 24 h 48 h
I.
Total Aurka (RT-qPCR)
Relative RNA Expression
J.
SUIT2
NTC SC 12 h 24 h 48 h
FC: 1.0 1.3 1.5 0.7 0.2
FC: 1.0 0.9 1.5 0.7 0.3
FC: 1.0 0.4 1.3 0.6 0.1
Western blot
K.
SUIT2
NTC SC 6 A
Radioactive RT-PCR
L.
+ Exon 2 Isoforms
PSI
NTC SC ASO-6 ASO-A
M.
Total Aurka (RT-qPCR)
Relative RNA Expression
N.
SUIT2
NTC SC 6 A
FC: 1.0 1.2 0.7 0.2
FC: 1.0 0.7 0.3 0.2
FC: 1.0 0.8 0.5 0.4
Western blot

Figure S10:

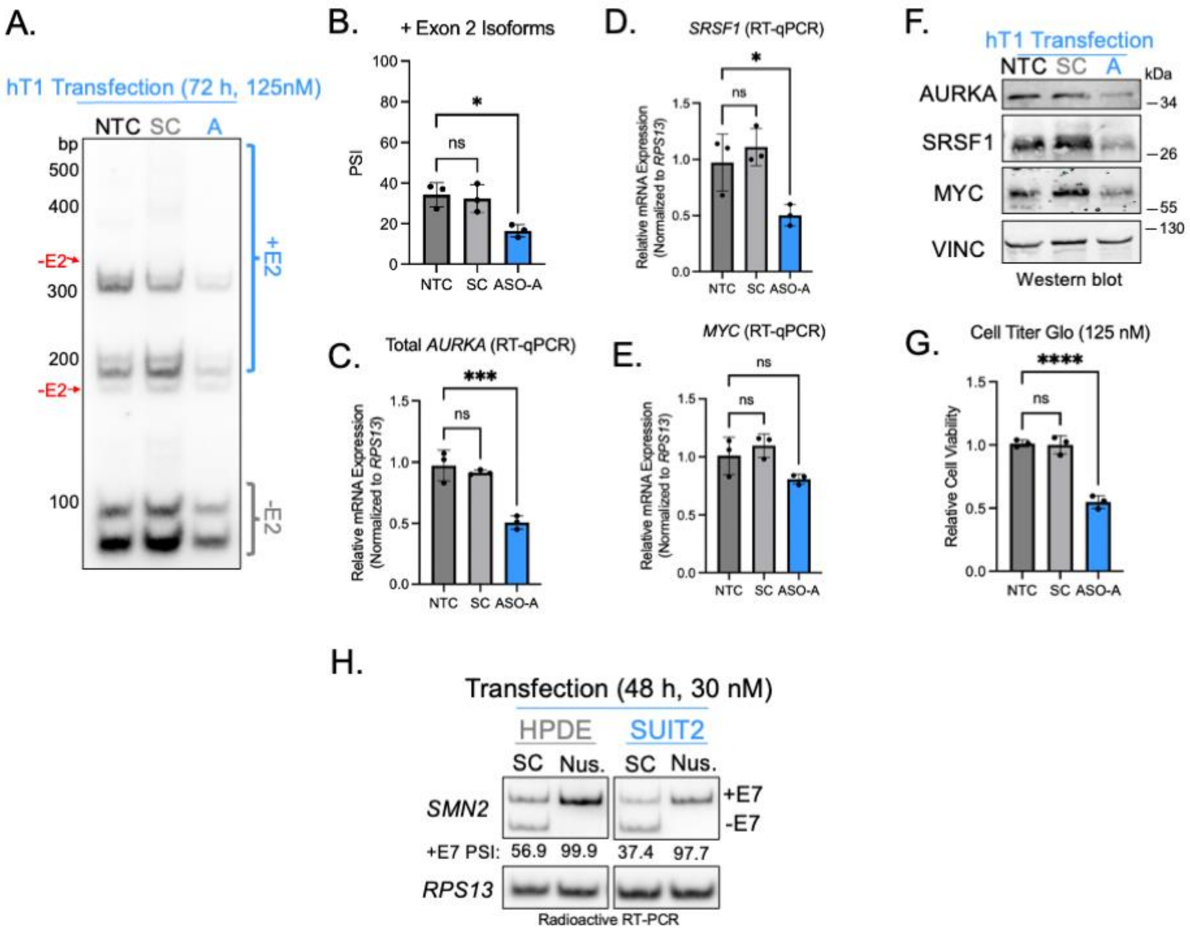
A.
hT1 Transfection (72 h, 125nM)
NTC SC A
bp
500
400
-E2
300
200
-E2
100
+E2
-E2
B.
+ Exon 2 Isoforms
PSI
ns
*
NTC SC ASO-A
C.
Total AURKA (RT-qPCR)
Relative mRNA Expression (Normalized to RPS13)
ns
***
NTC SC ASO-A
D.
SRSF1 (RT-qPCR)
Relative mRNA Expression (Normalized to RPS13)
ns
*
NTC SC ASO-A
E.
MYC (RT-qPCR)
Relative mRNA Expression (Normalized to RPS13)
ns
ns
NTC SC ASO-A
F.
hT1 Transfection
NTC SC A
kDa
AURKA
SRSF1
MYC
VINC
34
26
55
130
Western blot
G.
Cell Titer Glo (125 nM)
Relative Cell Viability
ns
****
NTC SC ASO-A
H.
Transfection (48 h, 30 nM)
HPDE
SUIT2
SC Nus. SC Nus.
SMN2
+E7
-E7
+E7 PSI: 56.9 99.9 37.4 97.7
RPS13
Radioactive RT-PCR

Figure S11:

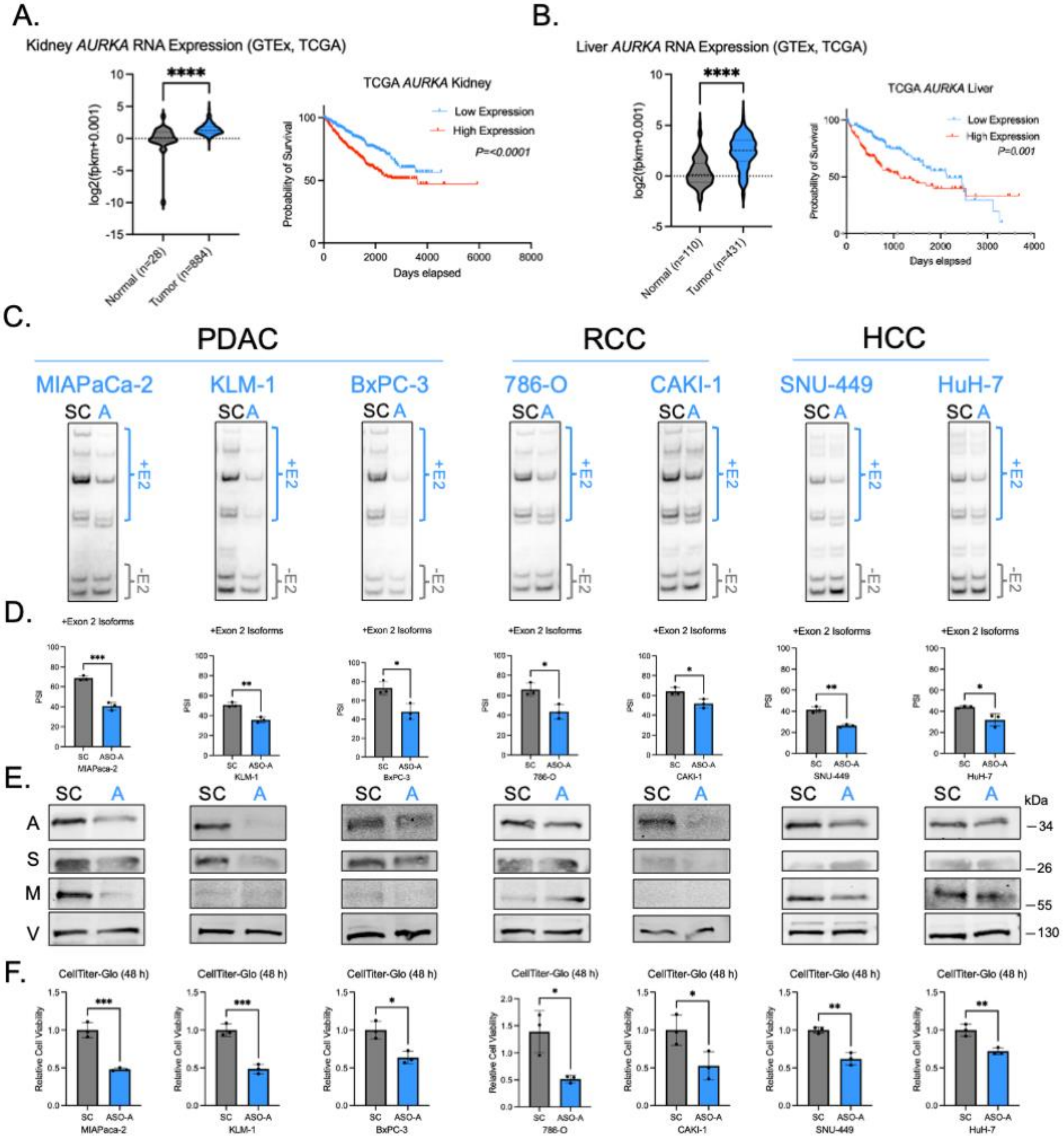

A.
Kidney AURKA RNA Expression (GTEx, TCGA)
TCGA AURKA Kidney
Low Expression
High Expression
P=<0.0001
B.
Liver AURKA RNA Expression (GTEx, TCGA)
TCGA AURKA Liver
P=0.001
C.
PDAC
RCC
HCC
MIAPaCa-2
KLM-1
BxPC-3
786-O
CAKI-1
SNU-449
HuH-7
D.
+Exon 2 Isoforms
E.
F.
CellTiter-Glo (48 h)

Table S1: Organoid Information

| **Organoid Line** | **Tissue Source** | **Tissue Type** | **Age at Consent** | **Birth Sex** | **Race** | **Class** | **KRAS** | **p53** |
|---|---|---|---|---|---|---|---|---|
| hT1 | Pancreas | Primary | 65 | Male | White | PDAC | G12V | R213 |
| hT60 | Pancreas | Primary | 57 | Male | NA | AMPULLARY | G12D | H193D |
| hT91 | Pancreas | Primary | 71 | Female | White | PDAC | G12D | T155N |
| hF3 | Pancreas | Primary | 69 | Female | Asian | PDAC | G12V | R273L |
| hF23 | Pancreas | Primary | 69 | Female | Black | PDAC | G12D | R175H |
| hM8 | Pancreas | Primary | 70 | Female | White | PDAC | G12R | LOSS |
| hM1A | Lung | Metastasis | 52 | Male | White | PDAC | G12D | R175H |
| hM19B | Liver | Metastasis | 43 | Male | White | PDAC | G12V | E298 |
| hN32 | Pancreas | Primary | 24 | Male | White | NORMAL | WT | na |
| hN33 | Pancreas | Primary | 28 | Male | White | NORMAL | WT | na |
| hN36 | Pancreas | Primary | 50 | Female | White | NORMAL | WT | na |
| hN39 | Pancreas | Primary | 53 | Male | Hispanic | NORMAL | WT | na |
| hN40 | Pancreas | Primary | 63 | Male | Hispanic | NORMAL | WT | na |

Table S2: Antisense oligonucleotides

| Name | Sequence | Chemistry |
|---|---|---|
| ASO-1 | AGACCCTGTCTCAAAAAAAA | * |
| ASO-2 | GAGTGAGACCCTGTCTCAAA | * |
| ASO-3 | CAATGGAGTGAGACCCTGTC | * |
| ASO-4 | CCCACCTCGGCCTCCGAAAA | * |
| ASO-5 | ATCCACCCACCTCGGCCTCC | * |
| ASO-6 | AGGTGATCCACCCACCTCGG | * |
| ASO-A | CTGGGCAATGGAGTGAGACC | * |
| ASO-3.2 | CTGGCCTGGGCAATGGAGTG | * |
| ASO-3.3 | GCACTCTGGCCTGGGCAATG | * |
| ASO-SC | GTCGCCTAAGCACTCACAGC | * |
| Nusinersen | ATTCACTTTCATAATGCTGG | * |

*All ASOs are uniformly modified with 2'-O-methoxyethyl (MOE) base modifications, phosphorothioate backbones, and all cytosines are methylated (5mC)

Table S3: PCR primers

| Name | Sequence (5'->3') | Experiments |
|---|---|---|
| SVP34_SRSF1_qPCR_F | CCGCAGGGAACAACGATTG | RT-qPCR |
| SVP35_SRSF1_qPCR_R | GCCGTATTTGTAGAACACGTCCT | RT-qPCR |
| SVP46_AURKA_splice_F | CTAACGGCTGAGCTCTTGGA | RT-PCR |
| SVP47_AURKA_splice_R | CAGGTCCTGAAATGCAGTTTTCT | RT-PCR |
| SVP55_AURKA_shRNA1_F | CCGGAGAATCCATTACCTGTAAATACTCGAGTATTTACAGGTAATGGATTCTTTTTG | Cloning |
| SVP56_AURKA_shRNA1_R | AATTCAAAAAGAATCCATTACCTGTAAATACTCGAGTATTTACAGGTAATGGATTCT | Cloning |
| SVP57_AURKA_shRNA2_F | CCGGCTGGCTCTTAAAGTGTTATTTCTCGAGAAATAACACTTTAAGAGCCAGTTTTTG | Cloning |
| SVP58_AURKA_shRNA2_R | AATTCAAAAACTGGCTCTTAAAGTGTTATTTCTCGAGAAATAACACTTTAAGAGCCAG | Cloning |
| SVP59_AURKA_qPCR_Total_F | GGAATATGCACCACTTGGAACA | RT-qPCR |
| SVP60_AURKA_qPCR_Total_R | TAAGACAGGGCATTTGCCAAT | RT-qPCR |
| SVP61_AURKA_qPCR_E2_F | GGTCTCACTCCATTGCCCAG | RT-qPCR |
| SVP62_AURKA_qPCR_E2_R | GAAAATGCTGGGATTACGGG | RT-qPCR |
| SVP63_mAurka_splice_F | GTAACGGCTGAGCTACCGGG | RT-PCR |
| SVP64_mAurka_splice_R | CAGGCCTGGAGACACAGTTTTCT | RT-PCR |
| SVP66_pGLO_Vec_Fwd | TGGGTCCTTGGGTCGCAGGTGCCACCATGGAAGATGCCAAAA | Cloning |
| SVP67_pGLO_Vec_Rev | GAGCTCAGCCGTTAGAATTCCGAAAGGCCCGGAGATGAG | Cloning |
| SVP68_pGLO_FRAG_FWD | CCTCATCTCCGGGCCTTTCGGAATTCTAACGGCTGAGCTCTTGG | Cloning |
| SVP69_pGLO_ISO1_Rev | TTGGCATCTTCCATGGTGGCACCTGCGACCCAAGGAC | Cloning |
| SVP70_pGLO_ISO2_REV | TTGGCATCTTCCATGGTGGCCTCGGCCTCCGAAAATGCT | Cloning |

| | | |
|---|---|---|
| SVP71_pGLO_ISO3_REV | TTGGCATCTTCCATGGTGGCCTGTAATCCCAGCTACTCGGGAG | Cloning |
| SVP72_pGLO_ISO4_REV | TTGGCATCTTCCATGGTGGCCTCGGCCTCCGAAAATGCT | Cloning |
| SVP73_pGLO_ISO5_REV | TTGGCATCTTCCATGGTGGCCTGTAATCCCAGCTACTCGGGAG | Cloning |
| SVP77_LUC2_qPCR_F | CCCATCTTCGGCAACCAGAT | RT-qPCR |
| SVP78_LUC2_qPCR_R | GTACATGAGCACGACCCGAA | RT-qPCR |
| SVP79_AURKA_ISO1_qPCR_F | GGGTCGCAGGCATCATG | RT-qPCR |
| SVP80_AURKA_ISO1_qPCR_R | CGAGAACACGTTTTGGACCTC | RT-qPCR |
| SVP81_AURKA_ISO2_qPCR_F | GGGTCGCAGGGTCTCACTC | RT-qPCR |
| SVP82_AURKA_ISO2_qPCR_R | GTCCATGATGCCTCGGCC | RT-qPCR |
| SVP83_AURKA_ISO3_qPCR_F | GTCGCAGGGTCTCACTCCA | RT-qPCR |
| SVP84_AURKA_ISO3_qPCR_R | ACACCATTGCACTCCAGCCTC | RT-qPCR |
| SVP85_AURKA_ISO4_qPCR_F | ATTACAGCTAGAGGGTCTCACTCC | RT-qPCR |
| SVP86_AURKA_ISO4_qPCR_R | CATGATGCCTCGGCCTC | RT-qPCR |
| SVP87_AURKA_ISO5_qPCR_F | ATTACAGCTAGAGGGTCTCACTCC | RT-qPCR |
| SVP88_AURKA_ISO5_qPCR_R | TTGCACTCCAGCCTCGG | RT-qPCR |
| SVP89_AURKA_ISO2a_qPCR_F | GTCGCAGGCTGGAGTGC | RT-qPCR |
| SVP90_AURKA_ISO2a_qPCR_R | CCATGATGCCTGTAATCCC | RT-qPCR |
| SVP99_SDMaurka_sh1res_fwd | CCTTGTCAGAATCCCTTGCCAGTGAATAGTGGCCAGG | Cloning |
| SVP100_SDMaurka_sh1res_R | CCTGGCCACTATTCACTGGCAAGGGATTCTGACAAGG | Cloning |
| SVP111_SDMaurka_sh2res_F | GTTTATTCTGGCTCTCAAGGTATTGTTTAAAGCTCAG | Cloning |
| SVP112_SDMaurka_sh2res_R | CTGAGCTTTAAACAATACCTTGAGAGCCAGAATAAAC | Cloning |
| SVP113_SDMaurka_D274N_F | CTTAAAATTGCAAATTTTGGGTGGTCAGT | Cloning |
| SVP114_SDMaurka_D274N_R | ACTGACCACCCAAAATTTGCAATTTTAAG | Cloning |
| SVP115_pCG_Vec_F | CAGCTAGCAAACAGTCTTAGCTCCTGGGCAACGTG | Cloning |
| SVP116_pCG_Vec_R | CCCTTGCTCACCATGGTGGCCCTGAAGTTCTCAGGATCCCC | Cloning |
| SVP117_pCG_AURKA_F | GGGATCCTGAGAACTTCAGGGCCACCATGGTGAGCAAGG | Cloning |
| SVP118_pCG_AURKA_R | ACCAGCACGTTGCCCAGGAGCTAAGACTGTTTGCTAGCTGATTCTTTGTT | Cloning |

| | | |
|---|---|---|
| SVP123_SRSF1_shRNA_F | CCGGACTGCCTACATCCGGGTTAAACTCGAGTTTAACCCGGATGTAGGCAGTTTTTG | Cloning |
| SVP124_SRSF1_shRNA_R | AATTCAAAAACTGCCTACATCCGGGTTAAACTCGAGTTTAACCCGGATGTAGGCAGT | Cloning |
| AK52_RPS13_F | CGAAAGCATCTTGAGAGGAACA | RT-qPCR |
| AK53_RPS13_R | TCGAGCCAAACGGTGAATC | RT-qPCR |
| SVP159_minigene_primer_F | GACCGAATCACCGACCTCTC | RT-PCR |
| SVP_minigene_primer_R | gcgctgggcccttcttaatg | RT-PCR |
| SVP221_SRSF1_Sh2_wan_F | CCGGGCAGAGGATCACCACGCTATTCTCGAGAATAGCGTGGTGATCCTCTGCTTTTTG | Cloning |
| SVP222_SRSF1_sh2_wan_R | AATTCAAAAAGCAGAGGATCACCACGCTATTCTCGAGAATAGCGTGGTGATCCTCTGC | Cloning |
| SVP264_MYC_qpcr_F | TCCCTCCACTCGGAAGGAC | RT-qPCR |
| SVP265_MYC_qpcr_R | CTGGTGCATTTTCGGTTGTTG | RT-qPCR |
| SVP272_Renilla_qpcr_F | GGGGTGCTTGTTTGGCATTT | RT-qPCR |
| SVP273_Renilla_qpcr_R | TCAGGCCATTCATCCCATGAT | RT-qPCR |
| SVP274_ASOres_muta_fwd | TTTGAGACAGGGTCTGACTGCATAGCCGAGGCCAGAGTGC | Cloning |
| SVP275_ASOres_muta_rev | GCACTCTGGCCTCGGCTATGCAGTCAGACCCTGTCTCAAA | Cloning |
| SVP308_MAGOH_shRNA1_F | CCGGCCAGAAGGCTTACGAGTATTTCTCGAGAAATACTCGTAAGCCTTCTGGTTTTTG | Cloning |
| SVP309_MAGOH_shRNA1_R | AATTCAAAAACCAGAAGGCTTACGAGTATTTCTCGAGAAATACTCGTAAGCCTTCTGG | Cloning |
| SVP310_MAGOH_shRNA2_F | CCGGCCGGACGGGAAGTTAAGATATCTCGAGATATCTTAACTTCCCGTCCGGTTTTTG | Cloning |
| SVP311_MAGOH_shRNA2_R | AATTCAAAAACCGGACGGGAAGTTAAGATATCTCGAGATATCTTAACTTCCCGTCCGG | Cloning |
| SVP312_Y14_shRNA1_F | CCGGCATCAGCGTTGACTGGTGTTTCTCGAGAAACACCAGTCAACGCTGATGTTTTTG | Cloning |
| SVP313_Y14_shRNA1_R | AATTCAAAAACATCAGCGTTGACTGGTGTTTCTCGAGAAACACCAGTCAACGCTGATG | Cloning |
| SVP314_Y14_shRNA2_F | CCGGCCAGGGATTATGATAGAAATACTCGAGTATTTCTATCATAATCCCTGGTTTTTG | Cloning |
| SVP315_Y14_shRNA2_R | AATTCAAAAACCAGGGATTATGATAGAAATACTCGAGTATTTCTATCATAATCCCTGG | Cloning |
| SVP322_MAGOH_qpcr_F | TTTATCTGCGTTACTACGTGGGG | RT-qPCR |
| SVP323_MAGOH_qPCR_R | CGTCCGGTCGAAACTCAAACT | RT-qPCR |
| SVP324_Y14_qPCR_F | GATGGGGACGAGAGCATTCAC | RT-qPCR |
| SVP325_Y14_qPCR_R | CGCTGTCATAATCCTCACGCA | RT-qPCR |
| SVP337_AURKA_3UTR_total_qPCR_F | ACCCCTGGATCACAGCAAAT | RT-qPCR |

| | | |
|---|---|---|
| SVP338_AURKA_3UTR_total_qPCR_R | AGCCCTGGCTCAAGGATTTC | RT-qPCR |
| SVP339_SRSF1_5UTR_total_qPCR_F | GGAAGGCCTGTTCTCGAGTC | RT-qPCR |
| SVP340_SRSF1_5UTR_total_qPCR_R | AGATGCGGCAATCGTTGTTC | RT-qPCR |
| SVP_AURKA_dI1_F | GGTCTCACTCCATTGCCCAGG | Cloning |
| SVP_AURKA_dI1_R | CTGCGACCCAAGGACCCAAG | Cloning |
| SVP_AURKA_dI2_F | GCTGGAGTGCAATGGTGTGATC | Cloning |
| SVP_AURKA_dI2_R | CTCGGCCTCCGAAAATGCTGG | Cloning |
| SVP_AURKA_dI3_F | GCATCCATGGAAGATGCCAAAAAC | Cloning |
| SVP_AURKA_dI3_R | CTGTAATCCCAGCTACTCGGGAGG | Cloning |
| YI390_Hs_MT16S_1258F | CCCTAGGGATAACAGCGCAA | RT-qPCR |
| YI391_Hs_MT16S_1359R | TAATAGCGGCTGCACCATCG | RT-qPCR |
| YI392_NEAT1_821F | ATTTTCCAGGTGGCAGTGCT | RT-qPCR |
| YI393_NEAT1_970R | CTTACTGTCCCGGGCTTACC | RT-qPCR |
| SVP_AURKA_3SS_F | AATGGTAATCTTAATTTTTTTTTTCTCTCAGGGTCTCACTCCATTGCCCAG | Cloning |
| SVP_3SS_R | ACCCTGAGAGAAAAAAAAAAATTAAGATTACCATTTTACAATTAAGCAAATAATTCCATGATGAAGCC | Cloning |
